# Phase Separation of a Novel form of Euchromatic Histone Methyltransferasel (EHMT1^N/C^) into cytoplasmic RNA viral Inclusion bodies facilitates their coalescence, thereby enhancing viral replication

**DOI:** 10.1101/2023.11.01.564969

**Authors:** Kriti Kestur Biligiri, Nishi Raj Sharma, Abhishek Mohanty, Debi Prasad Sarkar, Praveen Kumar Vemula, Shravanti Rampalli

## Abstract

Protein lysine methyltransferases (PKMTs) methylate histone and non-histone proteins to regulate biological outcomes such as development and disease including viral infection. While PKMTs have been extensively studied for modulating the antiviral responses via host gene regulation, their role in methylation of proteins encoded by viruses and its impact on host-pathogen interactions remain poorly understood. In this study, we discovered a distinct nucleo-cytoplasmic form of Euchromatic Histone Methyltransferase1(EHMT1^N/C^), a PKMT, that phase separates into viral inclusion bodies (IBs) upon cytoplasmic RNA-virus infection (Sendai Virus). EHMT1^N/C^ interacts with cytoplasmic EHMT2 and methylates SeV-Nucleoprotein upon infection. Elevated nucleoprotein methylation during infection correlated with coalescence of small IBs into large mature platforms for efficient replication. Inhibition of EHMT activity by pharmacological inhibitors or genetic depletion of EHMT1^N/C^ reduced the size of IBs with a concomitant reduction in replication. Since IB formation is conserved among all cytoplasmic RNA-viruses, our study will have strong implications in understanding the mechanisms regulating IB formation, discerning RNA viral pathogenesis and designing therapeutic strategies.

## Introduction

Euchromatic Histone N-lysine methyltransferase 1 (EHMT1) is an enzyme that belongs to the SET domain family of methyltransferases (1), known for its dual activity of methylating lysine residues (1) as well as binding methylated lysine residues (2). EHMT1, along with its homolog, EHMT2, forms heteromeric complexes (1,3) to methylate the ninth lysine residue of Histone 3 (H3K9), thereby transcriptionally repressing regions of the chromatin and signalling the formation of heterochromatin (1,4). The SET domain of EHMTs catalyse the transfer of methyl groups to lysine residues(1), and the ankyrin domain recognises and binds the methyl groups (2). In addition to histones, EHMTs methylate non-histone substrates to modulate a broad range of biological processes (5–9). Additionally, a high throughput study performed to determine the enzymatic substrates of EHMT1 and EHMT2 resulted in the identification of overlapping and non-overlapping substrates (6). Among these, several proteins were extranuclear, including several mitochondrial, ER and cell membrane specific proteins (6). While cytoplasmic isoform of EHMT2 is known (10–12), methylation of this vast array of EHMT1 specific extranuclear substrates couldn’t be explained as it was thought to be restricted to the nucleus(8).

The Histone methyltransferase activity of EHMTs has been extensively characterized in modulating the host antiviral response. For example, studies in embryonic stem cells identified that EHMT2 is required for exogenous retroviral silencing and MERVL ERVs (13,14). EHMT2 mediated H3K9me2 activity represses proinflammatory gene (IFN and IFN signalling genes) expression and upon viral infection this repression is relieved to resist the viral infection (15). The histone lysine methyltransferase activity of EHMT1 has also been characterised in modulating the host antiviral responses (15,16). For instance, EHMT1 maintains latency of retroviruses like HIV-1 by transcriptionally silencing the proviral elements integrated into the host genome (17). EHMT1 also regulates the expression of various inflammatory cytokines like IFNβ to maintain homeostasis under normal conditions (15,16). During infection by cytoplasmic RNA viruses like Influenza, Sendai and VSV, IFNβ expression is increased by releasing the repression laid by EHMT1 to inhibit viral replication(16). Overall, EHMTs are known to play an active epigenetic role during RNA viral infections by regulating the host antiviral response. However, the direct involvement of EHMTs in modulating RNA viral replication via its non-histone protein methylation has not been reported till date.

While performing cellular reprogramming experiments using Sendai virus (SeV) as a vector to deliver OSKM factors, we observed cytoplasmic condensation of EHMT1. Since EHMT1 is a nuclear protein, its unanticipated cytoplasmic localization led to further investigations, during which, we identified a novel nucleo-cytoplasmic form of EHMT1 (EHMT1^N/C^). We found that EHMT1^N/C^ acts as pro-viral host factor which was recruited to inclusion bodies of SeV. At mechanistic level, we discovered that cytoplasmic EHMTs (EHMT1^N/C^ and EHMT2) interact and methylate the nucleoprotein of SeV upon infection. Methyltransferase activity of EHMTs facilitated the formation of large replication platforms, thereby enhancing its propagation. Inhibition of EHMT’s enzymatic activity by small molecule inhibitors or reducing the levels of EHMT1^N/C^ by CRISPR/cas9 system revealed loss of larger IBs in infected cells indicating impairment in coalescence. Accordingly, we also noticed reduced replication of Sendai viral genomic RNA in infected cells that were inhibitor treated or gene edited. Overall, in this study we demonstrate for the first time, a novel nucleocytoplasmic form of EHMT1 along with cytoplasmic EHMT2, enzymatically promotes RNA viral replication via methylation of the nucleoprotein.

## Results

### Sendai virus induces cytoplasmic condensation of EHMT1

Ectopic expression of Yamanaka factors (OSKM) in somatic cells rewires the global epigenetic landscape towards pluripotent state (18–20). Such dramatic alterations are achieved by combinatorial efforts of several chromatin modifiers including EHMT1 (21,22). While the canonical role of EHMT1 has been studied during reprogramming, its non-canonical role remains poorly understood. Our study began with the intent to identify the non-canonical substrates of EHMT1 during the early phase of iPSC generation. Towards this, we adopted integration free [Cytotune-OSKM (Sendai viral vector) and episomal OSKM (plasmid based)] transgene delivery systems that are widely used in the field. We examined the pattern of expression of EHMT1 in 1. fibroblasts, 2. heterogenous cultures of OSKM transduced fibroblasts and in 3. iPSCs, by performing immunolabelling for EHMT1. Fibroblasts and iPSCs, demonstrated nuclear localization of EHMT1 (fig. S1A), which was consistent with its previously known sub-cellular localization (8). Interestingly, in the population of fibroblasts transduced with OSKM factors via Sendai Virus, we observed EHMT1 condensates of heterogenous size and shape distributed throughout the cytoplasm in addition to its expression in the nucleus (fig. S1A). Co­immunolabelling the heterogenous cultures with EHMT1 and Oct4 confirmed cytoplasmic condensation of EHMT1 in OSKM transduced cells (fig. S1B). However, surprisingly, such cytoplasmic condensates were completely absent in reprogramming cultures ectopically expressing OSKM via episomal plasmids (fig. S1C), indicating that the cytoplasmic localization of EHMT1 was not in response to OSKM expression. Since the localization of EHMT1 differed among the two modes of reprogramming, we speculated that its cytoplasmic condensation was not induced by reprogramming but was in response to the vector, Sendai virus. To test this, we infected fibroblasts with Cytotune EmGFP reporter SeV or Wild type (WT) SeV and performed immunolabelling for EHMT1. Cells infected with EmGFP SeV or WT SeV formed cytoplasmic condensates of EHMT1 of varying sizes and shapes (Fig.1, A and B) while retaining its nuclear localization, thus confirming our hypothesis. Additionally, we observed cytoplasmic EHMT1 condensation in other cell types like HEK, BEAS-2B and MEFs upon SeV infection (Fig.1, C and D, fig. S1D), indicating that our observations are not cell­type specific. Expression of Ezh2, another epigenetic modifier did not localize to cytoplasm in response to viral infection (Fig. 1E) indicating that this was not a common observation among all the epigenetic modifiers. Altogether, we identified that SeV infection induces cytoplasmic condensation of EHMT1, an epigenetic modifier, which was previously reported to be exclusively localized in the nucleus. Intrigued by the altered localization of EHMT1 in response to viral infection, we continued our investigation to uncover the role of EHMT1 in Sendai viral life cycle.

**Figure 1:**
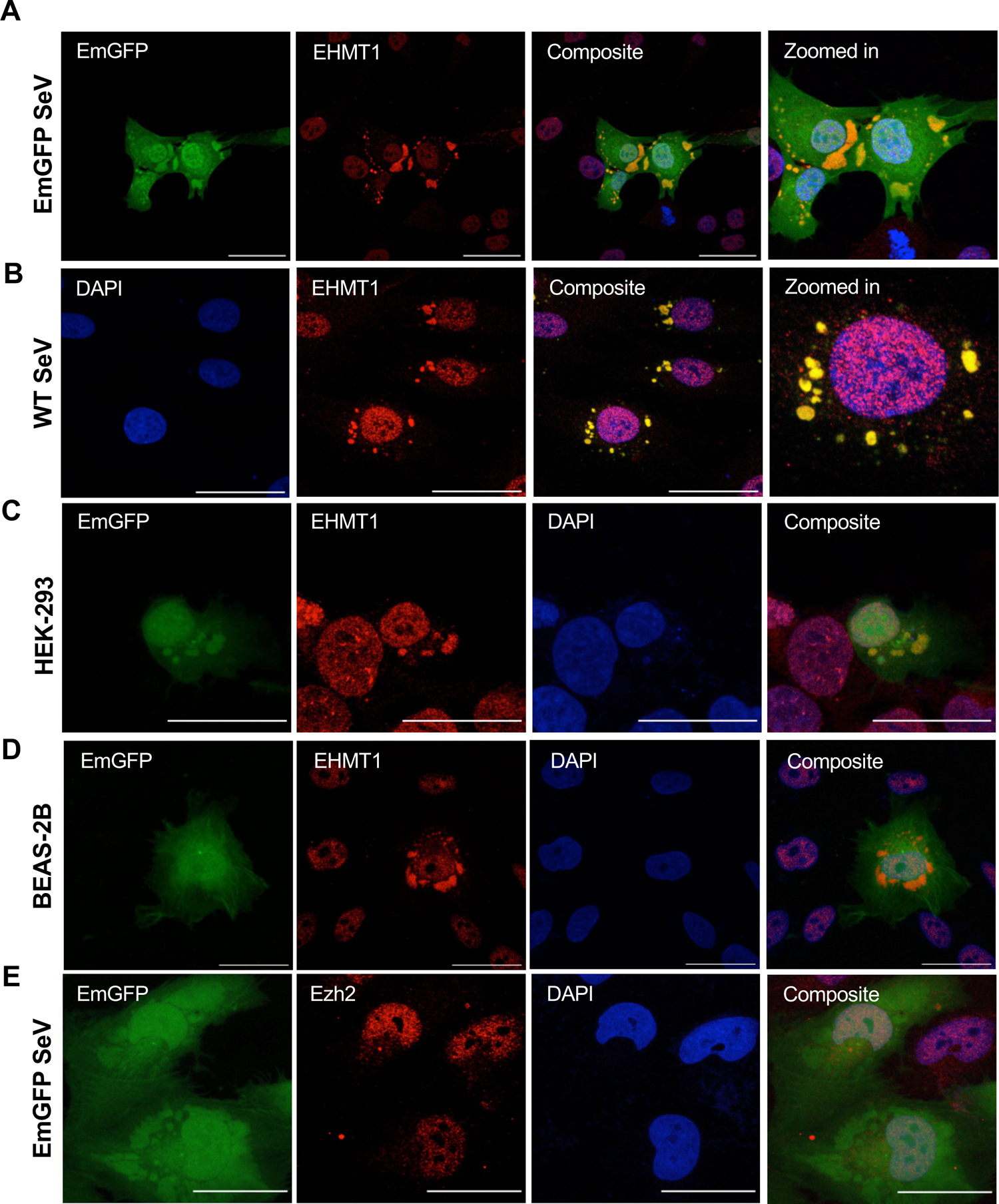
Sendai virus induces cytoplasmic condensation of EHMT1. Confocal microscopic images of (A) Human Dermal Fibroblasts (HDF) infected with EmGFP SeV (green), (B) HDF infected with WT SeV immunolabelled for EHMT1 (red). (C) HEK-293 cells infected with EmGFP SeV (green), immunolabelled for EHMT1 (red) (D) BEAS-2B cells infected with EmGFP SeV (green), immunolabelled for EHMT1 (red) or (E) Ezh2 (red) 48h p.i.. Composite images are with DAPI (blue) stained nuclei. Scale bar - 40pm.

### EHMT1 is recruited to Sendai viral Inclusion Bodies

Sendai virus, a member of the paramyxoviridae family of single strand negative sense RNA viruses, replicates in the cytoplasm of the host (23–25). Its genomic RNA is bound by the Nucleoprotein (N), a ssRNA binding protein, Phosphoprotein (P), a cofactor of the polymerase, and the RNA dependent RNA Polymerase (or L) to constitute an RNP complex. Upon entering the host, the viral RNP complex of most members of the paramyxoviridae family serves as a template for transcription (26–29). As the concentration of N, P and L proteins increase, they tend to phase separate from the aqueous cytosol, forming subcellular structures known as Inclusion Bodies (IBs) (26–29). By restricting the exchange of material between IBs and cytosol, these structures protect the viral components from being detected and degraded by the host immune surveillance machinery (30,31). N and P proteins also recruit several host factors to IBs (30–35), thus accumulating the necessary viral and host components within these subcellular structures for viral replication. While the role of viral components in IB formation has been studied, the host factors recruited to IBs and their contribution to IB formation and viral replication remains to be determined.

IBs formed by members of the paramyxoviridae family like measles virus are cytoplasmic condensates of irregular shapes and sizes. Since EHMT1 formed similar condensates in the cytoplasm in response to SeV infection, we examined if these are SeV IBs. EmGFP SeV or WT SeV infected cells were co­immunolabelled with EHMT1 and a polyclonal SeV antibody, where we found that EHMT1 colocalised with SeV (Fig.2, A and B), indicating that SeV proteins and EHMT1 localise to the same sub-cellular compartment. To determine if these structures are indeed IBs, we used SeV specific markers like the N and P proteins, which were cloned into an mCherry_C1 plasmid (mCh_N) and piRFP_N1 plasmid (piRFP_P) respectively. Transfection of mCh_N resulted in diffused cytoplasmic expression of N and EHMT1 (fig. S2, A and B) but upon infection, cytoplasmic condensates of mCh_N were observed (Fig. 2C), marking the IBs. ICC against EHMT1 demonstrated a colocalization of EHMT1 with SeV N in viral condensate (Fig. 2C). Similarly, piRFP_P when transfected independently, was diffused in the cytoplasm (fig. S2, C and D) but condensed into SeV IBs upon infection, where EHMT1 was found to localise (Fig. 2D). Thus, in SeV infected cells, EHMT1 condensed into viral IBs, marked by the N and P proteins. In addition to the viral markers of IBs, certain host proteins like Hsp70 have been reported to interact with the viral replication proteins of Mumps (36), Rabies (26,37) and Respiratory Syncytial virus (38,39) in their IBs, where they directly influence viral replication. We then examined if Hsp70 is recruited to SeV IBs by co-immunolabelling for Hsp70 and EHMT1 in SeV infected cells, where we found that Hsp70 localised to SeV IBs (fig. S2, E and F), which are marked by N, P and EHMT1 proteins.

**Figure 2:**
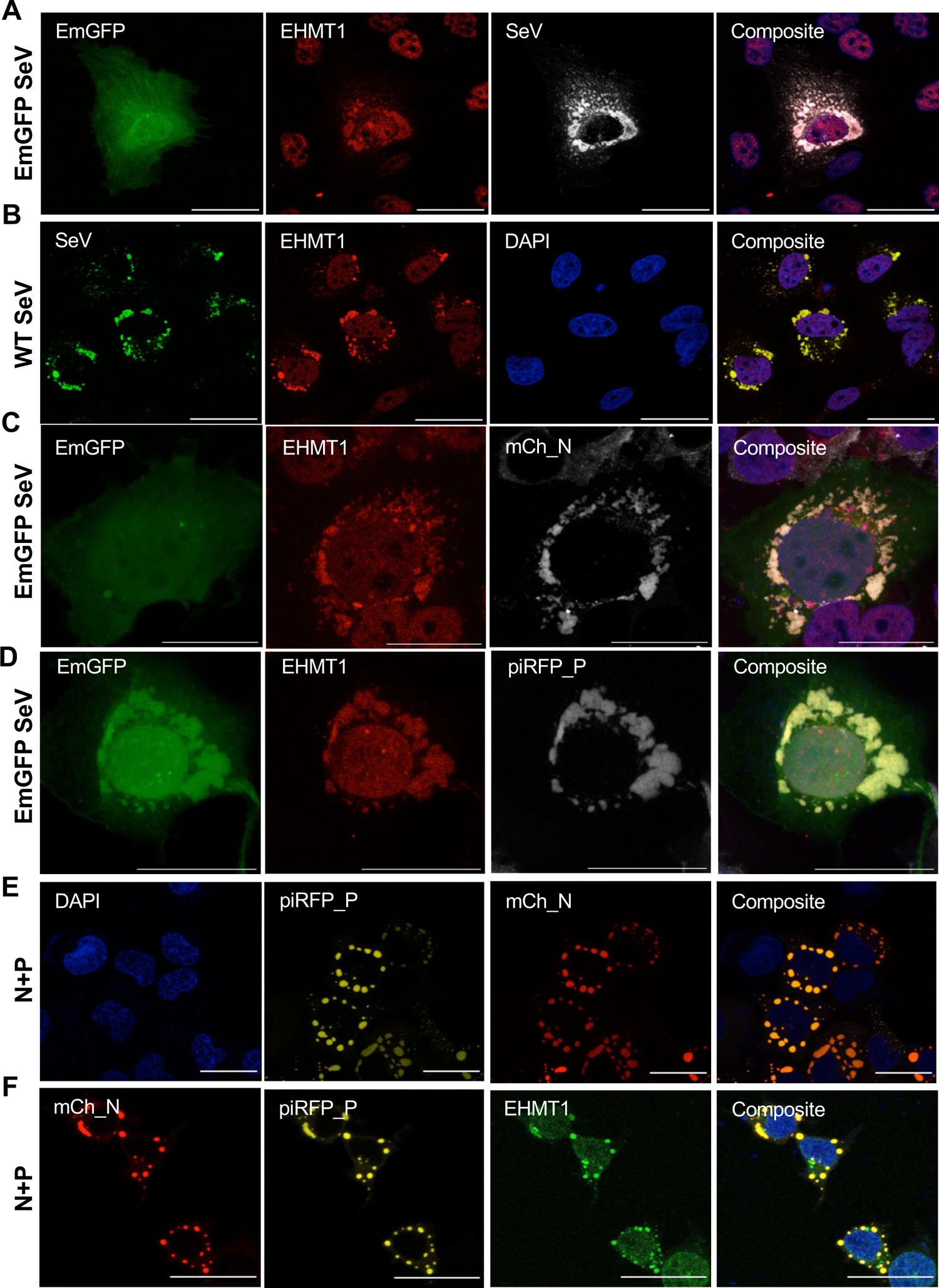
EHMT1 is recruited to Sendai viral Inclusion Bodies. Confocal microscopic images of (A) BEAS-2B infected with EmGFP SeV (green) co-immunolabelled with EHMT1 (red) and SeV (gray), (B) BEAS-2B infected with WT SeV co-immunolabelled with EHMT1 (red) and SeV (green). (C) HEKs infected with EmGFP SeV (green) and transfected with mCh_N (gray), immunolabelled with EHMT1 (red), (D) HEKs infected with EmGFP SeV (green), transfected with piRFP_P (gray) and immunolabelled with EHMT1 (red). (E) HEK co-transfected with mCh_N (red) and piRFP_P (yellow) at a ratio of 1:1 (F) HEK co-transfected with mCh_N (red) and piRFP_P (yellow) at a ratio of 1:1 and immunolabelled with EHMT1 (green). Composites of all images are with DAPI (blue) stained nuclei. Scale bar - 40μm.

The N and P proteins of viruses belonging to the order mononegavirales form phase separated compartments in the cytosol upon their co-expression, independent of infection (28,29,40). Although the IBs formed by co-transfection of N and P are inert structures as opposed to the IBs formed during infection, these IBs mimic the structures formed upon infection in terms of the factors recruited and their properties (28–32,40). SeV N and P proteins upon co-transfection also formed spherical shaped IBs (Fig. 2E), resembling liquid phase separated structures. ICC against Hsp70 in co-transfected cells revealed a localization of Hsp70 with the IBs (fig. S2G), indicating that the recruitment of certain host proteins is conserved with the IBs formed during infection. We next examined if N and P are responsible and sufficient for the recruitment of EHMT1 in the absence of infection, towards which, we performed ICC against EHMT1 in co-transfected cells. We found that EHMT1 localised to the IBs formed by N and P independent of viral RNA (Fig. 2F). Altogether, we identified that EHMT1 is a part of SeV IBs, which are marked by the viral N and P proteins and a host protein, Hsp70. The expression of replication proteins, N and P suggests that these IBs serve as replication sites. Additionally, we demonstrated that N and P proteins are the factors responsible for the condensation of EHMT1 into IBs.

### A distinct nucleo-cytoplasmic form of EHMT1 associates with SeV IBs

Since the time EHMT1 was identified, it has been known to localise exclusively in the nucleus. In this study, for the first time, we observed extranuclear localization of EHMT1 into cytoplasmic SeV IBs. Considering that viral IBs recruit host proteins (30–39), we hypothesised that SeV could be inducing nuclear to cytoplasmic shuttling of EHMT1 upon infection. To test our hypothesis, we cloned the full length EHMT1 in a mCherry_C1 vector (mCh_EH1_FL) (fig. S3A). Transfection of mCh_EH1_FL followed by confocal microscopic imaging demonstrated its localization to the nucleus (fig. S3B). We then infected the mCh_EH1_FL transfected cells with SeV to examine its cytoplasmic shuttling. Surprisingly, we did not observe any cytoplasmic mCh_EH1_FL signal in response to SeV infection (fig. S3C). While SeV IBs were formed in the cytoplasm, mCh_EH1_FL expression was restricted to the nucleus of infected cells (fig. S3C), indicating that the full-length nuclear EHMT1 doesn’t shuttle, and the cytoplasmic protein might be different from the nuclear form.

EHMT1 and its homolog, EHMT2, express several isoforms, among which, EHMT2 is known to express a cytoplasmic isoform (10–12). Thus, we hypothesised that a unique cytoplasmic isoform of EHMT1 might be expressed in response to SeV infection. To test this hypothesis, we fractionated SeV infected and uninfected cells into the cytoplasmic and nuclear fractions. As seen in Fig. 3A, the nuclear fraction was enriched with LaminB1 and cytoplasmic with Gapdh. To our surprise, we observed EHMT1 in the cytoplasmic fraction irrespective of SeV infection and the cytoplasmic EHMT1 resolved slightly lower than nuclear EHMT1 (Fig. 3A). This indicated the expression of two forms of EHMT1, a full-length form in the nucleus and a slightly smaller form in the cytoplasm. Since these two forms had only a minute size difference, we could not segregate them in the whole cell lysate. To sort this, we resolved the fractions on a lower percentage (4%) SDS-PAGE gel, which revealed that smaller form of EHMT1 was not unique to the cytoplasm but was also expressed in the nucleus (Fig. 3B). We termed this isoform of EHMT1 as EHMT1^N/C^. Quantification of the levels of EHMT1 in the nuclear and cytoplasmic fractions revealed no significant increase in either of the compartments upon infection. (Fig. 3A and C, fig S3D). However, the levels of both nuclear and cytoplasmic EHMT1 reduced slightly upon infection (Fig. 3, A and C, fig S3D), which was consistent with a global quantitative proteomic study performed on cells infected with Sendai Virus (41).

**Figure 3:**
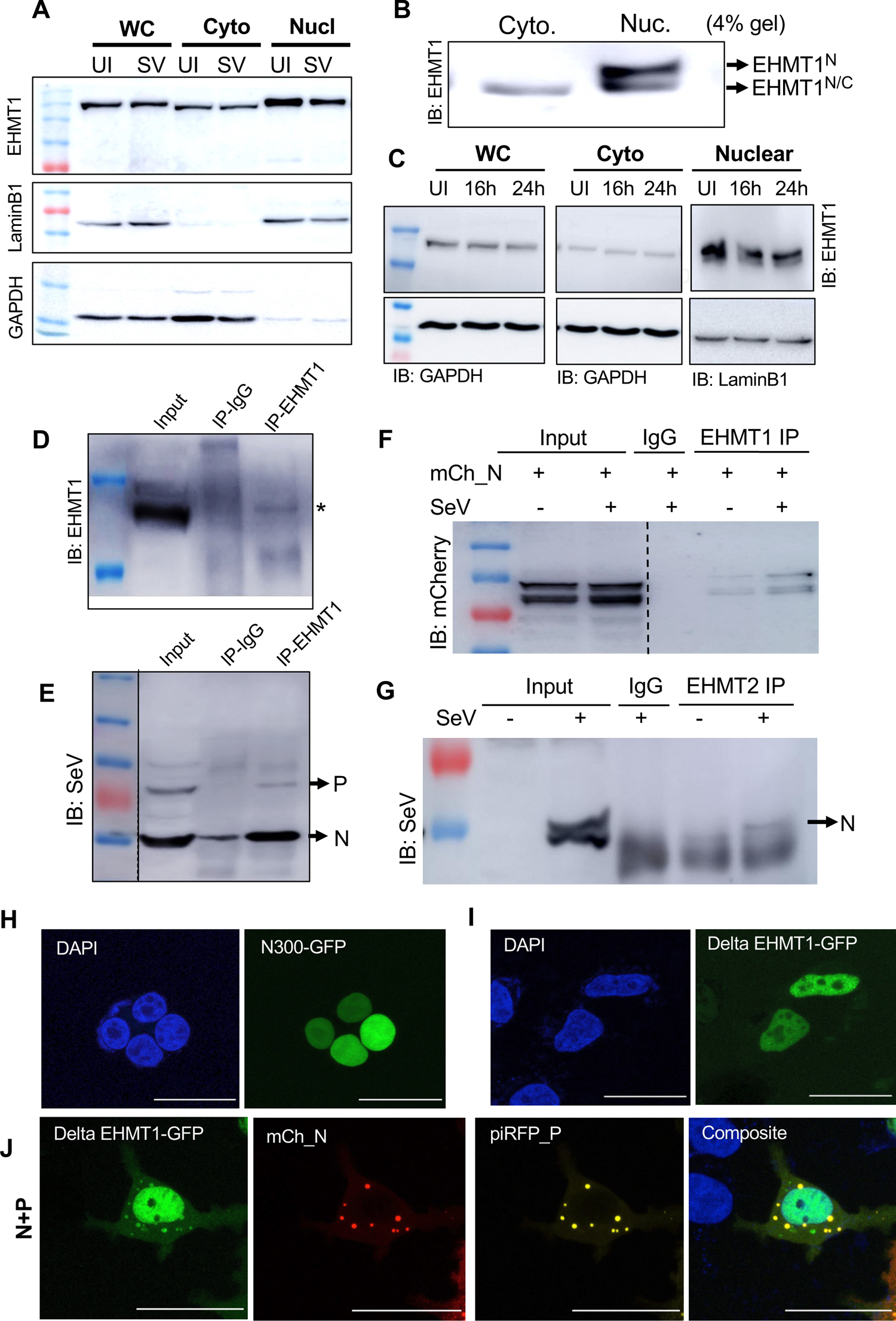
A distinct nucleo-cytoplasmic form of EHMT1 associates with SeV IBs. (A) Western Blotting of the whole cell, nuclear and cytoplasmic fractions of Uninfected BEAS-2B cells (UI) and EmGFP SeV infected (SV) at 48h p.i.. Blots probed with EHMT1, LaminB1 and Gapdh. (B) Nuclear and cytoplasmic fractions of BEAS-2B cells resolved on a 4% SDS-PAGE gel, immunoblotted with EHMT1. (C) WB of nuclear and cytoplamic fractions of BEAS-2B cells uninfected and infected with WT SeV at 16h and 24h p.i., probed with EHMT1, GAPDH and LaminB1. (D) and (E) EHMT1 IP performed on the cytoplasmic fraction of SeV infected cells, elute Western Blotted and probed with EHMT1 (Upper panel) and SeV (Lower panel). (F) EHMT1 IP from the cytoplasmic fraction of cells transfected with mCh_N, with and without SeV infection, elute Western Blotted and probed with mCherry. (G) EHMT2 IP from the cytoplasmic fraction of SeV infected and uninfected cells, elute Western Blotted and probed with SeV. Confocal microscopic images of HEK transfected with (H) N300-GFP (green), (I) Delta-EHMT1-GFP (green), (J) Delta-EHMT1-GFP (green), co-transfected with mCh_N (red) and piRFP_P (yellow). Composites of all images are with DAPI (blue) stained nuclei. Scale bar - 40μm.

To examine if EHMT1^N/C^ associates with viral proteins, we immunoprecipitated EHMT1 from the cytoplasmic fraction of SeV infected cells and immunoblotted for SeV and EHMT1. Our data revealed an association of EHMT1^N/C^ (Fig. 3D) with two viral proteins, SeV-N (57 KDa) and SeV-P (62 KDa) (42) (Fig. 3E), of which, there was a stronger enrichment of N than P. Our data thus indicated that a distinct cytoplasmic form of EHMT1, EHMT1^N/C^, gets recruited to SeV IBs, where it associates with two key viral proteins critical for viral replication and IB formation. To conclusively determine whether EHMT1 indeed interacts with the Nucleoprotein, we transfected cells with the SeV nucleoprotein tagged with mCherry (mCh_N), which was followed by SeV infection. EHMT1 was then immunoprecipitated from the cytoplasmic fraction of cells transfected with mCh_N and infected with SeV. The elute was then resolved by Western blotting and probed for mCherry, where we could detect that EHMT1 indeed associated with the Nucleoprotein (Fig. 3F). Further, to ensure that cytoplasmic EHMT1 is indeed the protein interacting with SeV proteins, we transfected cells with the nuclear mCh_EH1_FL and infected with SeV. Immunoprecipitation of mCherry from the whole cell lysate of these cells revealed no association with SeV proteins, indicating that the cytoplasmic and not the nuclear EHMT1 associates with SeV proteins (fig. S3E).

EHMT1 and EHMT2 are known to form functional heterodimers; since EHMT2 has a cytoplasmic isoform, we examined for any possible interaction between the two proteins in the cytoplasm. Towards this, we immunoprecipitated EHMT1 from the cytoplasmic fraction of uninfected and infected cells and found an association between the two proteins in the cytoplasm (fig. S3F). Further, we also immunoprecipitated EHMT2 from the cytoplasm of SeV infected cells and found an association with the SeV nucleoprotein (Fig. 3G). Thus, we found that cytoplasmic EHMT1 and EHMT2 associate with each other as well as the viral Nucleoprotein upon infection.

Compared to EHMT1_N_, EHMT1^N/C^ exhibited a small size difference, such variations are usually a resultant of PTMs like phosphorylation or due to the formation of a novel transcript with deletion of small exons from the full-length mRNA. Alternative splicing of the exon10 of EHMT2 results in an isoform that shuttles between the nucleus and cytoplasm (10,11). Given this precedence, we first examined if there is loss of a portion of EHMT1 which can confer the differential localization. EHMT1 protein can be broadly divided into two halves (1-735 and 735 - 1296). Second half consists of the Ank and catalytic SET domain while the first half is majorly comprised of the disordered domains. To date, there is no information on the NLS or NES signals of EHMT1, however, there are predicted monopartite and bipartite sequences present in exons 5, 7, 8, 13 and 14. To find out the potential sequence whose loss might alter the localization of FL-EHMT1, we generated 3 constructs N300 (Exon 1-5), ΔEHMT1 (Exon 5-13 deleted)) and Ank+SET (Exon 13-27) (fig. S3G). The constructs upon transfection, revealed that the Ank+SET domains remained in the cytoplasm (fig. S3H) and its overexpression formed aggregation in the cytoplasm (fig. S3H). While both N300 and ΔEHMT1 showed nuclear localization (Fig. 3, H and I), we also observed that in few cells exclusively in ΔEHMT1 transfected, a faint GFP signal was seen in the cytoplasm (Fig. 3I). Intrigued by this observation, we co-transfected N300 and ΔEHMT1 transfected cells with SeV nucleoprotein and phosphoprotein. To our surprise, ΔEHMT1 colocalised with SeV IBs (Fig. 3J) but such change in localization was not seen in N300 alone (fig. S3I). These data indicated that loss of a small region between exon 5-13 might impart properties of distinct subcellular localization to EHMT1. Nonetheless, we identified a novel short form of EHMT1 which is present in both the nucleus and the cytoplasm and associates with SeV IBs.

### EHMTs regulate SeV IB formation and viral replication

EHMTs are lysine methyltransferases known to methylate histone and several non-histone proteins, thereby regulating biological process. To examine the requirement of EHMT’s enzymatic activity in viral processes such as IB formation and replication, we inhibited the methyltransferase activity of EHMTs using small molecule inhibitors BIX01294 (43) and UNC0642 (44). BEAS-2B cells were treated with 3μM BIX or 3μM UNC and infected simultaneously with WT SeV, where IB formation was examined by immunolabelling for SeV at 16h p.i. (Fig. 4A). In untreated cells, SeV IBs were compact structures of heterogenous sizes ranging from small to large, whereas in BIX and UNC treated cells, the IBs were predominantly smaller in size (Fig. 4B), indicating that EHMTs might have a role to play in the formation of large IBs. We then assessed the impact of EHMTs inhibition on replication of SeV, where we found a 30 - 40% reduction in viral gRNA replication upon inhibition (Fig. 4C). Similarly, reduction in the number and size of IBs were noticed upon inhibition of EHMTs in cells infected with recombinant EmGFP SeV (Fig. 4D and E). Further, replication of the EmGFP SeV was also reduced by about 50% (Fig. 4F) at 24h p.i..

**Figure 4:**
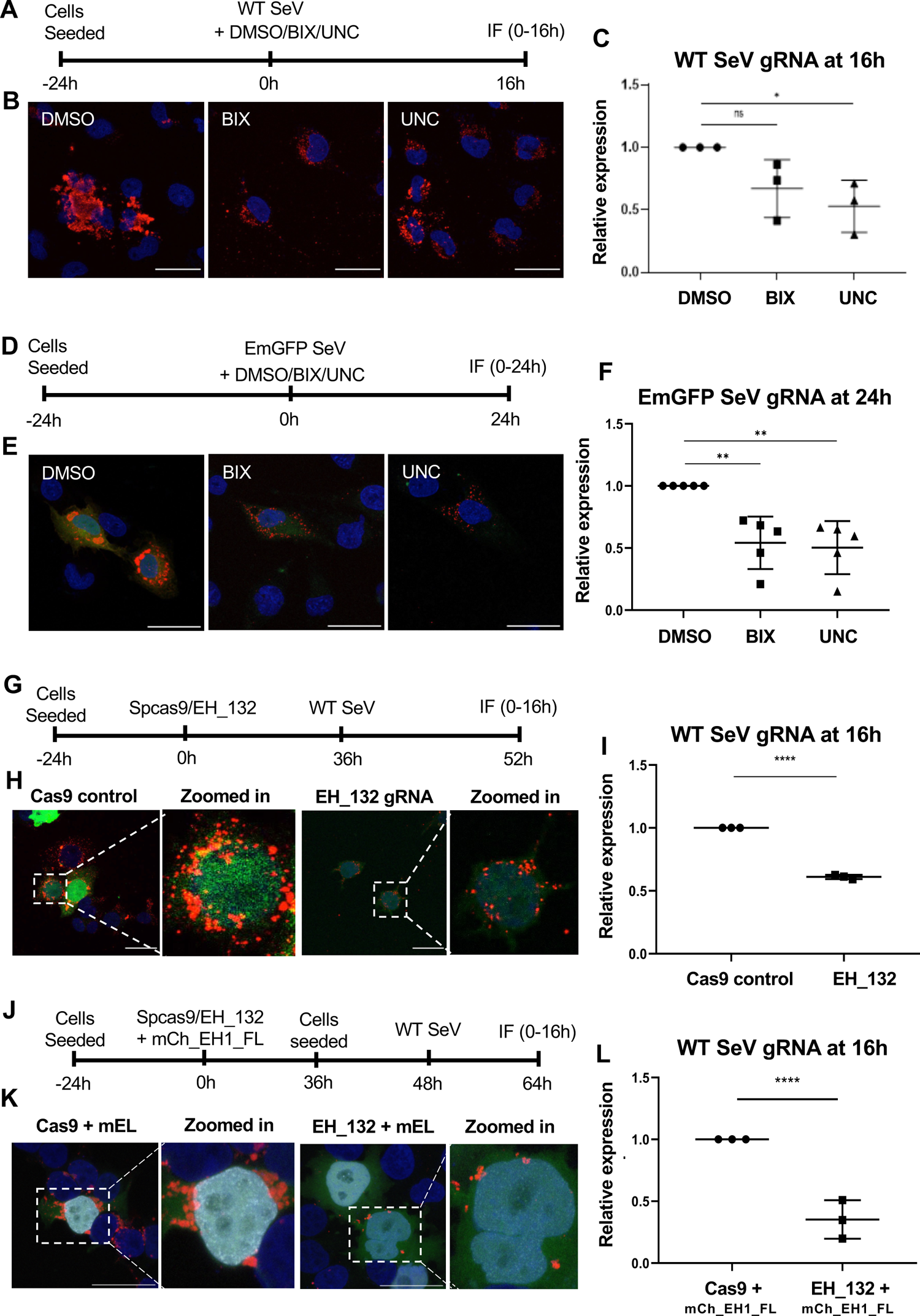
EHMTs regulate SeV IB formation and viral replication. (A) Experimental scheme representing BEAS-2B cells treated at 0h with DMSO, 3μM BIX or 3μM UNC and infected with WT SeV (B) Confocal microscopic composite images of cells immunolabelled with SeV ab (red), marking the IBs. (C) Graphs plotted for the relative expression of SeV gRNA assessed by qRT-PCR with the Ct values normalised against GAPDH. (n=3 replicates, one-way ANOVA, **, p<0.005, *, p<0.05, ns, p=0.1129) (D) Experimental scheme representing BEAS-2B cells treated at 0h with DMSO, 3gM BIX or 3gM UNC and infected with EmGFP SeV. (E) Confocal microscopic composite images of EmGFP SeV infected cells (green), immunolabelled with SeV ab (red), marking the IBs (F) Graphs plotted for the relative expression of SeV gRNA assessed by qRT-PCR with the Ct values normalised against GAPDH (n=3 replicates, one-way ANOVA, **, p<0.005, *, p<0.05, ns, p=0.1129). (G) Schematic representation of the experimental procedure. (H) Confocal microscopic composite images of the cells transfected with empty cas9 plasmid or EH_132 (green reporter), immunolabelled for SeV ab (red). (I) Graph plotted for the relative expression of SeV gRNA normalised with GAPDH as assessed by qRT-PCR. (J) Schematic representation of the experimental procedure. (K) Confocal microscopic composite images of cells transfected with spcas9/EH_132 (green), mCh_EH1_FL (gray) and immunolabelled with SeV ab (red). Nuclei of all images are counterstained with DAPI (blue). Scale bar - 40μm. (L) Graph plotted for the relative expression of SeV gRNA normalised with GAPDH as assessed by qRT-PCR. (Data from I and L represent mean ± S.D. (n=3 replicates, unpaired t-test, ****, p<0.0001).

Observation of EHMT1 in this cellular compartment, where its expression hadn’t been reported previously led us to employ an alternative approach to specifically verify if EHMT1^N/C^ is required for IB formation and viral replication. We depleted the levels of endogenous EHMT1 by using the CRISPR/cas9 system. A guide RNA targeting the exon3 of EHMT1 (fig. S4A) was cloned into pspcas9(BB)-2A-GFP plasmid (EH_132), which was transfected in HEK to deplete EHMT1. The target site of the EH_132 was validated for mutation by T7 endonuclease assay (fig. S4B). We obtained about 40 - 60% reduction in the protein levels of EHMT1 (fig. S4C) in cells transfected with EH_132, where we also observed a reduction in the H3K9me2 levels (fig. S4C). We further sorted the GFP positive cells to obtain clonal populations. These cells were fractionated into the nuclear and cytoplasmic fractions respectively, to determine the levels of EHMT1_N_ and EHMT1^N/C^. The protein was found to be significantly depleted in both the nuclear as well as the cytoplasmic fraction (fig. S4D). Cells that exhibited complete loss of EHMT1, proliferated extremely slowly and could not be propagated to establish a knockout line. Thus, we proceeded with HEK demonstrating about 70% depletion of EHMT1, infected them with WT SeV 36h post transfection, where we immunolabelled for SeV and imaged the infected cells positive for the GFP reporter to assess IB formation (Fig. 4G). We observed that the size of IBs were reduced (Fig. 4H), which aligned with our observations upon inhibition of its methyltransferase activity. Assessing the SeV gRNA levels in EHMT1 depleted cells by qPCR indicated a reduction of replication by about 40%, indicating that the recruitment of EHMT1 to SeV IBs facilitates their formation, which in turn supports viral replication (Fig. 4I).

In the above experiments, EH_132 gRNA depleted both the nuclear and cytoplasmic form of EHMT1, therefore it was difficult to assess whether the effects observed on IB formation was solely in response to the loss of cytoplasmic form of EHMT1. To dissect the roles of nuclear vs cytoplasmic EHMT1, we devised an experimental setup in which we depleted both the nuclear and cytoplasmic forms of EHMT1 by CRISPR/cas9. Additionally, we selectively introduced the nuclear full length EHMT1 in EHMT1 depleted cells. This resulted in a system where the nuclear EHMT1’s expression was restored but the cytoplasmic EHMT1 was depleted. Towards this, we transfected EH_132 transfected cells with EHMT1 full length construct, tagged with mCherry (mCh_EHMT 1 _FL) to compensate for the loss of the nuclear protein (fig. S4E). These cells were then infected with WT SeV to specifically determine the effects of loss of cytoplasmic EHMT1^N/C^ on viral replication (Fig. 4J). Immunocytochemistry of SeV infected cells that were expressing gRNA (Cas9 or EH_132) and mCh_EHMT1_FL identified that the nuclear full length EHMT1_N_ could not rescue the defective IB formation (Fig. 4K). While the nuclear EHMTl’s activity has been implicated in regulating an antiviral response elicited by the host (15–17), our data demonstrate that a cascade reaction instigated by nuclear EHMT1 via epigenetic mechanisms may not implicated in cytoplasmic IB formation. This data further reiterated the role of the cytoplasmic EHMT1^N/C^ in formation of IBs. Consequently, we observed about 50% reduction in viral gRNA replication in cytoplasmic EHMT1 depleted cells (Fig. 4L). Overall, our results thus demonstrated that both inhibition of the enzymatic activity as well as depletion of the protein resulted in disrupted IB formation and reduction in viral replication.

### Incorporation of EHMT1^N/C^ into SeV IBs correlates with the formation of large IB platforms and enhanced SeV replication

For cytoplasmic RNA viruses, IB formation is a dynamic process wherein small IBs coalesce with each other to form large structures to enhance viral replication (45,46). Since the process of IB formation hasn’t been studied for Sendai virus, we wanted to determine the mechanism of IB formation and the contribution of EHMT1 towards formation of IBs. To address this, we monitored the behaviour of SeV IBs using live cell microscopy in HEKs transfected with mCh_N and infected with EmGFP-SeV (Fig. 5A and movie S1). IBs formed in infected cells exhibited heterogeneity in size and shape; monitoring them over the time course of a few minutes led to the observation that spherical shaped small and intermediate sized IBs were mobile and appeared to coalesce to form larger structures (Fig. 5A and movie S1); indicating that coalescence of smaller IBs might give rise to large IBs. Additionally, IBs also underwent fission (Fig. 5A and movie S1), indicating a possibility of crosstalk among neighbouring IBs and the environment, a phenomenon reported to occur in IBs of several members of the order mononegavirales (27,28,46,47). Thus, SeV IBs, like other members of the family, exhibited properties of fusion and fission.

**Figure 5:**
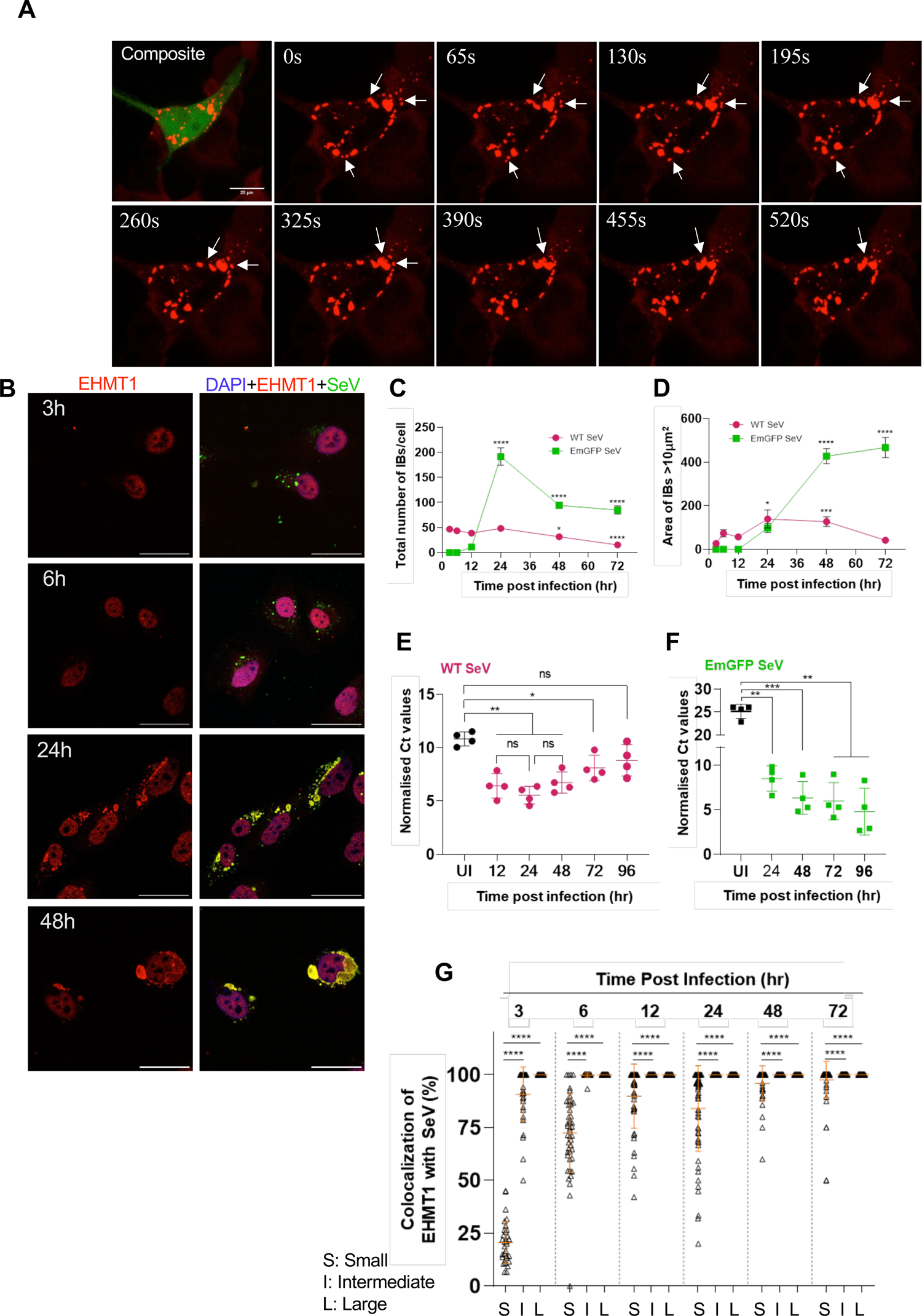
Incorporation of EHMT1^N/C^ into SeV IBs correlates with the formation of large IB platforms and enhanced SeV replication. (A) Live cell confocal microscopic images of HEK transfected with mCh_N (red) and infected with EmGFP SeV (green) at 48h p.i.; arrows represent fusion and fission of SeV IBs. (B) Confocal microscopic images of BEAS-2B cells infected with WT SeV, fixed and co-immunolabelled for EHMT1 (red) and SeV (green) at indicated time points. (C) and (D) BEAS-2B cells infected with WT or EmGFP SeV were fixed and immunolabelled with SeV at various time points post infection. Confocal microscopic images were analysed by ImageJ. (C) Graph representing the ± S.E.M. of total number of IBs formed per cell (pink line - WT SeV, green line - EmGFP SeV), (D) Graph representing the area ± S.E.M. of IBs >10μm^2^ per cell (pink line - WT SeV green line - EmGFP SeV) [n>25 cells, Brown-Forsythe and Welch ANOVA tests]. (E) and (F) Replication of SeV genomic RNA at indicated time points post infection. Graphs representing the Ct values for SeV genomic RNA normalised with GAPDH for (E) WT SeV and (F) EmGFP SeV. Data from E and F are mean ± S.D. (n=4 replicates, Repeated Measures (RM) One-way ANOVA). (G) BEAS-2B cells were infected with WT SeV, fixed and co-immunolabelled with EHMT1 and SeV at indicated time points post infection. Graph representing the percentage colocalization of EHMT1 with SeV in the three subpopulations of IBs, analysed by the Volocity image analysis software. Data are ± S.E.M. (n>25 cells, Ordinary One-way ANOVA). p-value: 0.1234 (ns), 0.0332 (*), 0.0021 (**), 0.0002 (***), <0.0001 (****)).

Next, we studied the dynamics of IB formation by analysing the number and size of IBs formed at various stages of infection. Towards this, we infected BEAS-2B cells with either WT or EmGFP SeV and co-immunolabelled with EHMT1 and SeV at various time points post infection. In WT SeV infected cells, IBs were detected as early as 3h p.i. throughout the cytoplasm, where they appeared as tiny spherical speckles (Fig. 5B). As the infection progressed, most of the spherical speckle-like IBs transformed into larger structures with irregular shape (Fig. 5B). To gain clarity into the formation of IBs, we first quantified the total number of IBs formed per cell from 3h to 72h post infection with WT SeV (Fig. 5C). We observed that the total numbers remained constant from 3h to 12h p.i., peaked at 24h, followed by a steep decline post 24h, indicating that 24h could be the point of peak infection (Fig. 5C). Our data on the total number of IBs was a resultant of all IBs irrespective of their heterogeneity, which was observed among infected cells at given time points as infection progressed.

Studies performed on parainfluenza and metapneumovirus revealed that formation of large IBs is crucial for efficient viral replication (45,46). To understand the dynamics of SeV IB formation and its relationship with replication, we first measured the size of each IB at various time points. Quantifying the size of IBs revealed that IBs ranged from 0.1μm^2^ to >100pm^2^, which led us to categorise them into four classes based on their size: Small (0.1-1μm^2^), intermediate (1-10pm^2^), large (10-100pm^2^) and very large (>100 pm^2^). We observed that small IBs dominated the population from 3h to 24h, post which, their numbers declined at 48h and 72h (fig. S5A). Intermediate IBs were the second most dominant population in the above mentioned timepoints and demonstrated a similar trend of decline (fig. S5A). Large IBs were very few in numbers, which remained constant till 12h, however, their population doubled by 24h (Fig. 5D, fig. S5A), post which, there was a steep decline in their numbers. Additionally, we also observed the emergence of very large IBs at 24h and 48h (Fig. 5D, fig. S5A). Overall, this data indicated that while small and intermediate IBs increased in numbers slightly to attain a peak at 24h, followed by their decline, number of the large IBs doubled by 24h and very large IBs emerged at 24h and 48h. A decline in the number of small/intermediate IBs at the time point of emergence of large/very large IBs indicated coalescence of smaller IBs to form larger structures. To understand if the formation of larger IBs correlated with SeV replication, we quantified viral genomic RNA over the indicated time points (Fig. 5E). A peak in SeV gRNA replication was observed at 24h (Fig. 5E), at which point, the number of large and very large IBs peaked, implying that the formation of large IBs correlated with higher replication.

For EmGFP SeV, we observed an initial delay in IB formation, where the earliest IBs were formed at 12h p.i. (Fig. 5C and fig. S5B), followed by a rapid increase in their numbers by 24h, with steep decline at 48h and 72h. IBs at 12h appeared as tiny spherical speckles (fig. S5B), which increased in numbers by 24h (Fig. 5C), contributing to a major portion of the IB population. This was followed by intermediate and large IBs, very large IBs were occasionally observed in cells at later time points of infection. While the number of small, intermediate, and large IBs peaked at 24h (fig. S5A), followed by a steep decline from 48h onwards, we observed a peak in the number of very large IBs at 48h and 72h (Fig. 5D, fig. S5A and B). This was like the trend observed with WT SeV IBs, where an increase in the number of very large IBs coincided with the point of decline of smaller IBs, indicating coalescence of smaller IBs to form larger structures as infection progressed. Quantitative analysis of the viral genomic RNA (Fig. 5F) further confirmed a peak in replication at the time points of formation of very large IBs. Collectively, these data indicated an increase in the size of IBs as their numbers decreased over the time course of infection. Additionally, the appearance of very large IBs correlated with an increase in viral gRNA replication, confirming that coalescence of IBs to form large structures is essential for efficient replication.

Since inhibition/depletion of EHMT1 led to the formation of smaller IBs (Fig. 4, B, E, H and K), we examined if a relationship existed between the size of IBs and the inclusion of EHMT1^N/C^. To examine this, we assessed the co-localization of EHMT1^N/C^ with SeV in IBs of different sizes at various time points post infection. Object-based colocalization analysis was performed on confocal microscopic images using the Volocity Image Analysis software, where IBs were marked by SeV antibody and immunolabelled with EHMT1^N/C^. At the earliest time point of WT SeV infection (3h), we observed that about 10% of small, 85% of intermediate and 100% large/very large IBs contained EHMT1^N/C^ (Fig. 5G). As the infection progressed, there was a gradual but substantial increase in incorporation of EHMT1^N/C^ into small IBs with an increase from 10% at 3h to about 80% at 72h (Fig. 5G). Intermediate IBs exhibited about 100% colocalization by 12h whereas, large/very large IBs exhibited 100% incorporation throughout (Fig. 5G). This data demonstrated that EHMT1^N/C^ is incorporated in all the large IBs; its incorporation in small/intermediate IBs increased as the infection progresses, thus demonstrating a clear relationship between the size of IBs formed during infection and the incorporation of EHMT1^N/C^. Altogether, our data indicated that SeV IBs exhibit properties of fusion and fission and formed large IBs by coalescence. In addition to this, a strong correlation exists between the size of IBs and incorporation of EHMT1^N/C^, indicating a possible role of EHMT1^N/C^ in promoting maturation or formation of SeV IBs to generate larger platforms for efficient replication.

### EHMTs methylate the SeV Nucleoprotein upon infection

Recent studies have unravelled that IBs are sites of viral genomic RNA replication where complexes of the N and P have been known to facilitate viral replication (26,27,34,45–47). Like other members of the paramyxoviridae family, complexes of the N, P and L have been identified to facilitate SeV replication as well (23). Since EHMTs are associated with N and P proteins, we speculated it to be a component of the viral RNP complex. To examine if EHMT1^N/C^ is indeed recruited to sites of SeV gRNA replication, we immunoprecipitated EHMT1^N/C^ from the cytoplasmic fraction of SeV infected cells, followed by qPCR of the immunoprecipitated elute to amplify SeV gRNA. We observed a 1000­fold enrichment of SeV gRNA in EHMT1 IP compared to beta-actin, which was used as a negative control (Fig. 6A), indicating that EHMT1 is a part of the RNP complex.

**Figure 6:**
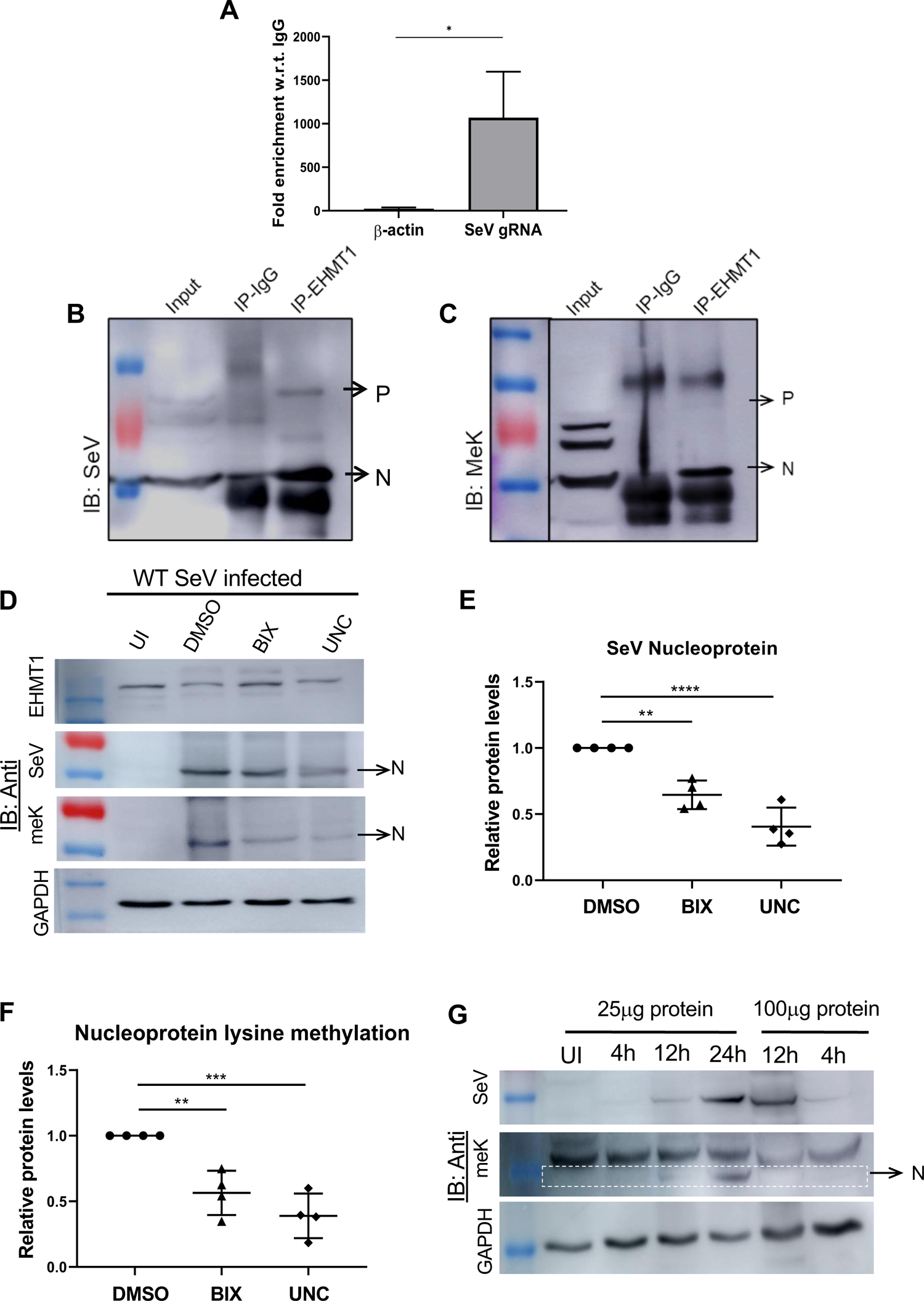
EHMTs methylate the SeV Nucleoprotein upon infection. (A) EHMT1 RNA-IP graph representing the fold enrichment of beta-actin and SeV gRNA, normalised with IgG. Data are mean ± S.D. (n=3, Ratio paired t-test; *, p<0.05). SeV gRNA demonstrated about 1000-fold higher enrichment in comparison with beta-actin. (B) and (C) BEAS-2B cells were infected with EmGFP SeV and EHMT1 was immunoprecipitated from the cytoplasmic fraction 48h p.i.. IP elute was Western blotted and probed with (B) SeV and (C) meK. (D) BEAS-2B cells were treated with BIX/UNC and simultaneously infected with WT SeV, cells were fractionated 24h post infection. The cytoplasmic fraction was Western blotted and probed with EHMT1, SeV, meK and GAPDH. (E) and (F) Graphs representing quantification of the SeV nucleoprotein (E) and (F) meK levels from Western blots. For (E), protein levels were normalised against GAPDH, for (F), Protein levels were normalised against the respective nucleoprotein bands. (n=4 replicates, one-way ANOVA, **, p<0.005, 0.0002 (***), <0.0001 (****)). (G) Western blotting of lysates from UI, 4h, 12h and 24h post infection of HEK with WT SeV, blots probed with SeV, meK and GAPDH.

Nucleo and phospho proteins, the two key components of the SeV RNP complex, associates with and recruits EHMT1^N/C^ to IBs (Fig. 2F and 3E), where its methyltransferase activity was found to facilitate the formation of large IBs (Fig. 4, B, E, H and K). These observations prompted us to speculate whether SeV N and P proteins could be the enzymatic substrates of EHMTs to promote the formation of large IBs. Thus, to decipher this, we examined the status of methylation of the N and P proteins in SeV infected cells. Towards this, we immunoprecipitated EHMT1 from the cytoplasmic fraction of SeV infected cells, one half of the elute was immunoblotted with SeV antibody (Fig. 6B), where we found an association of EHMT1 with the SeV N and P proteins. The other half of the elute probed with meK antibody (Fig. 6C) revealed methylation of the Nucleoprotein and Phosphoprotein. The N and P proteins, which are components of the viral RNP complex, have several overlapping functions in the viral life cycle such as regulation of IB formation and viral replication (Figure 3). Since EHMT was found to regulate these stages of the viral life cycle, we decided to probe into the activity of EHMT on one of its potential substrates, the Nucleoprotein, which is also the most abundantly expressed viral protein.

To specifically determine whether EHMT is the lysine methyltransferase responsible for methylating the SeV Nucleoprotein, we treated the cells with EHMT specific inhibitors BIX and UNC while simultaneously infecting them with WT SeV. The cells were fractionated 24h p.i. to obtain the cytoplasmic fraction, which was Western blotted to assess the levels of the Nucleoprotein, which revealed a decrease in the levels of the Nucleoprotein (Fig. 6D) by about 40 - 50% (Fig. 6E). We have reported that inhibition of EHMT1 resulted in reduction in size of the viral IBs, which in turn reflected on a reduction in viral RNA replication (Figure 3). This effect probably translates to the downstream processes like viral protein synthesis, which explains a reduction in the levels of the Nucleoprotein.

Therefore, to assess the effect of EHMT on methylation of the Nucleoprotein, we parallelly probed the blot with methylated lysine (meK) antibody, where we observed a reduction in its levels upon EHMT inhibition (Fig. 6D). To specifically determine the effect of EHMT on methylation of the Nucleoprotein while negating the effects of the inhibitors on the reduced viral replication, we quantified the meK bands by normalizing them with the respective Nucleoprotein bands. This led to the finding that the lysine methylation of the Nucleoprotein was further reduced by about 50% in EHMT inhibitor treated cells (Fig. 6F) irrespective of the overall reduced levels of the Nucleoprotein. This data thus indicated that EHMTs are the specific methyltransferases responsible for lysine methylation of the Nucleoprotein.

In Fig. 5G, we reported time dependent recruitment of EHMT1 into SeV IBs and its direct corelation with the replication of SeV and size of the IBs. Since the recruitment of EHMT1 was 100% in large IBs, whose population seemed to be affected by the inhibition/depletion of EHMT1, we questioned if this reflected on the methylation of Nucleoprotein. To address this, we harvested WT SeV infected cells at various time points (4h, 12h and 24h) post infection, wherein we observed a gradual increase in the population of large IBs (Fig. 5, B and D; fig S5A). Since 4h is very early during infection, we detected weak expression of Nucleoprotein by Western Blotting and the amount of N increased gradually from 12h to 24h (Fig. 6G). Probing the blot with meK antibody revealed an increase in methylation of N from 12 to 24h (Fig. 6G). However, this observation could be attributed to an overall increase in N in a time dependent manner. To circumvent this technical concern, we used 4 times the amount of 12h lysate for normalization with 24h, which gave us nearly equal amounts of N at both the time points (Fig. 6G). Interestingly, although the levels of nucleoprotein were equal in both time points, the meK levels were very low at 12h compared to 24h (Fig. 6G), indicating that methylation of nucleoprotein is a time dependent phenomenon. Thus, an incremental recruitment of EHMT1 to SeV IBs over the time course of infection is reflected on its methyltransferase activity on the SeV nucleoprotein.

Next, we wanted to test if infection is the determining factor for methylation of the Nucleoprotein by EHMT1. So, we used the N and P co-transfection system, where IBs formed independent of viral infection. Examining the association of EHMT1 with N and P demonstrated a weak association with the nucleoprotein (fig. S6A and B). However, we did not detect methylation of Nucleoprotein in the absence of infection (fig. S6C). These findings indicated that EHMT1 neither strongly associates with nor does it methylate the nucleoprotein independent of infection. Furthermore, quantitation of the size of IBs in N and P co-transfected cells revealed that they were comprised of small and intermediate sized IBs (fig. S6D). We rarely observed large IB in these cells, whereas very-large IBs were completely absent. These results were further supported by the fact that IBs formed by N and P co-transfection did not attain the size of IBs formed during infection (fig. S6D), indicating the possible role of Nucleoprotein methylation in formation of large IBs.

### Methyltransferase activity of EHMT1^N/C^ catalyses the growth of IBs to form larger structures

To examine if EHMTs indeed promote the maturation/growth of SeV IBs, we studied the effect of inhibition/depletion of EHMT1 on the various subpopulations of IBs. Towards this, we treated BEAS- 2B cells with small molecule inhibitors of EHMT1 & 2, BIX and UNC and simultaneously infected them with WT SeV. We performed ICC against SeV to mark the IBs to determine the impact of EHMT’s inhibition on their formation. Analysis of the total number of IBs formed per cell (Fig. 7A) demonstrated about 20% increase in the number of IBs in inhibitor treated cells. Measuring the overall size distribution of the whole IB population (Fig. 7B) led to the finding that the size of IBs in untreated cells ranged from 0.1 μm^2^- 1000 μm^2^, which was reduced to 0.1μm^2^ - 90pm^2^ upon EHMT1 & 2 inhibition. To understand the relationship between an overall increase in the number of total IBs and a reduction in their mean size, we studied the subpopulations of IBs. The number of small IBs demonstrated about 37% increase in BIX and 24% increase in UNC treated conditions (fig. S7A) whereas intermediate IBs treated with BIX demonstrated a decrease and UNC demonstrated a slight but significant increase in numbers (fig. S7B). However, the number of large IBs declined by 77% in BIX and 70% UNC treated cells (fig. S7C), indicating that this is the most affected population. Thus, an increase in the number of small IBs and decrease in the number of large IBs upon inhibition of the enzymatic activity of EHMT1^N/C^ indicated an inhibition of coalescence. To assess the extent of inhibition of IB growth, we measured the size of large IBs (>10pm^2^) (Fig. 7C), where we found that DMSO treated cells formed IBs with a mean area of greater than 100pm^2^ whereas, the mean area in inhibitor treated cells was about 20pm^2^ (Fig. 7C), indicating about 82% reduction in BIX and 75% reduction in UNC treated conditions. Thus, this data clearly indicated that EHMTs catalyse the coalescence and subsequently the growth of small IBs to form large structures.

**Figure 7:**
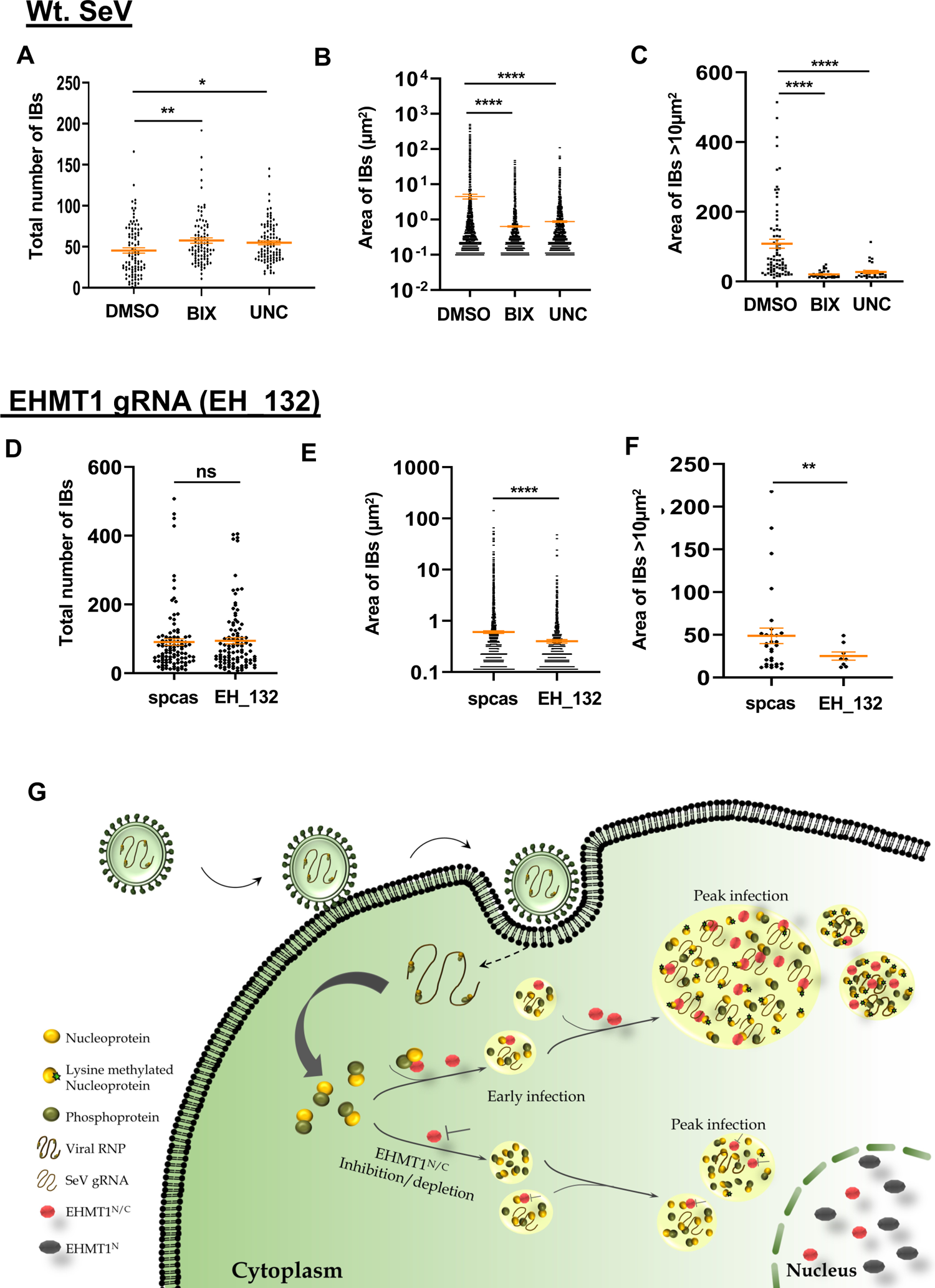
Methyltransferase activity of EHMT1^N/C^ catalyses the growth of IBs to form larger structures. (A) to (C) BEAS-2B cells were infected with WT SeV and treated with 3μm of BIX/UNC. The cells were fixed and immunolabelled with SeV antibody 16h p.i., the number of IBs and their area was quantified by ImageJ. Graphs representing (A) Total number of IBs per cell, (B) Distribution of area of the whole IB population, (C) Area of IBs >10μm^2^. (N=3 replicates, n>100 cells, Ordinary one­way ANOVA). (D) to (F) HEK293 cells were transfected with EH_132 plasmid to deplete the levels of EHMT1 and infected with WT SeV, cells were immunolabelled with SeV 16h p.i., number and area of IBs was quantified by ImageJ. Graphs representing (D) Total number of IBs per cell, (E) Distribution of area of the whole IB population, (F) Area of IBs >10μm^2^. Data from A to F are ± S.E.M (N=3 replicates, n>100 cells, Unpaired t-test), p-value: 0.1234 (ns), 0.0332 (*), 0.0021 (**), 0.0002 (***), <0.0001 (****). (G) Model figure: Upon SeV infection, the cytoplasmic form of EHMT1, EHMT1^N/C^ is recruited to SeV IBs by the N and P proteins in a time dependent manner as infection progresses. EHMT1^N/C^ and cytoplasmic EHMT2 plays an enzymatically active role in coalescence of IBs by methylating N, leading to the formation of large IBs and efficient viral replication. Inhibition of EHMTs activity or depletion of EHMT1^N/C^ leads to impairment in coalescence of IBs.

Next, we performed similar analysis in EHMT1 depleted cells, infected with WT SeV. Towards which, CRISPR/cas9 targeting of EHMT1 was employed; EH_132 plasmid (with GFP reporter) was transfected in HEK to deplete the levels of EHMT1. GFP positive infected cells were imaged from both spcas9 empty vector and EH_132 transfected cells. By analysing the IBs marked by SeV antibody at 16h p.i., we observed a small increase in the total number of IBs (Fig. 7D) and a minute decline in the mean size of IBs (Fig. 7E) formed upon depletion of EHMT1. Upon classification of IBs into their subpopulations, the number of small IBs were found to increase by 10% (fig. S7D) whereas intermediate IBs displayed a 52% reduction in numbers (fig. S7E) and large IBs reduced by 76% (fig. S7F) in EHMT1 depleted population. Analysing the size of large IBs formed revealed about 60% decrease in the size of the largest IB formed upon depletion of EHMT1 (Fig. 7F). We believe that the subtle effects on the increase in small IBs and decrease in large IBs compared to that of the inhibitor could be due to about 40% EHMT1 present in the system (fig. S4C).

Since EH_132 targeted both nuclear and cytoplasmic EHMT1, we wanted to specifically determine the effects of cytoplasmic EHMT1 on IB formation. To address this, we transfected EH_132 transfected cells with mCh_EHMT1_FL to compensate for the activity of the nuclear EHMT1. These cells were then infected with the WT SeV and immunolabelled with SeV antibody to assess IB formation. Quantification of the IBs thus formed revealed no significant increase in the total number of IBs (fig. S7G) but a decrease by about 50% in the area of IBs >10um2 (fig. S7H). These results aligned with our observations on cells depleted of total EHMT1, thus indicating that the effect of EHMT1 observed on IB formation is a resultant of the cytoplasmic EHMT1^N/C^.

Further, to convincingly demonstrate the requirement of EHMT1^N/C^ in the formation of large IBs for efficient SeV replication, we studied IBs formed independent of infection. Since EHMT1^N/C^ was also recruited to IBs formed by the co-expression of SeV N and P proteins in absence of other viral components, we assessed the effects of its inhibition on the IBs thus formed. The total number of such IBs per cell were very few in comparison with those formed upon SeV infection; the size of these IBs were also smaller, where they ranged from 0.1-10 pm^2^ (small and intermediate IBs) (fig. S7I). Very rarely, we noticed the formation of large IBs in these co-transfected cells. Treatment with BIX or UNC didn’t impact the number or size of these IBs (fig. S7I), implying that the role of EHMT1^N/C^ is confined to the formation of large IBs during viral replication. Altogether, our data on EHMT1^N/C^ incorporation into IBs (Fig. 5G), its enzymatic inhibition and depletion (Fig. 4, A to I) convincingly demonstrate that the methyltransferase activity of EHMT1^N/C^ is essential for formation of large IBs, which is critical for efficient replication (Fig. 7G).

## Discussion

RNA viruses are obligate parasites which require the host cellular machinery to complete their life cycle. These viruses trigger profound changes in the gene expression of specific host proteins and bring about global reorganization of the host cell proteome to facilitate their replication (38–40,48). Notably, infection by these viruses induce the formation of biomolecular condensates called inclusion bodies (26–29). Depending on the stage of infection, the size, shape and number of IBs varies in an infected cell (28,29,45). Formation of IBs is initiated by aggregation of newly synthesized IDR-rich viral nucleo of phospho proteins (28,40). During early phase, IBs are small with minimal viral proteins and their size increases as the infection progresses. Larger IBs represent mature IBs which have been shown to contain host factors including chaperones (26,30–34,36–39,48,49). Given that different steps of viral life cycle occur within IBs, the biogenesis and maturation of these structures are crucial for viral replication (49,50). Understanding these processes may lead to understanding pathogenesis of these viruses and designing therapeutic interventions.

Formation of such distinct cytoplasmic organelles is regulated by the concentration of proteins and nucleic acids, their composition, chaperones and PTMs (51–56). While the concentration and composition of biomolecules has been studied by in vitro assays, deciphering how PTMs influence IBs has been challenging due to the technical difficulties involved in isolating these structures sensitive to ionic strength and detergents (56–59). Nonetheless, few PTMs like SUMOylation, arginine methylation, phosphorylation and ubiquitylation have been reported to directly influence biomolecular condensates but a vast majority of them remains to be studied (57,58,60–64). In this regard, our study identified a protein lysine methyltransferase, EHMT1^N/C^ as a pro-viral host factor for SeV pathogenesis and provides insights into how SeV engages with EHMT1^N/C^ for successful replication. We report that upon SeV infection, EHMT1^N/C^ localizes to viral cytoplasmic condensates as early as 3h. Among the six SeV proteins, EHMT1^N/C^ associates with the nucleoprotein and phosphoproteins and their co-expression in cells was sufficient for the condensation and recruitment of EHMT1^N/C^. At the functional level, localization of EHMT1^N/C^ into viral IBs lead to formation of larger IBs and efficient SeV replication. Consistently, we observed small IBs and reduced replication upon knockdown or inhibition of EHMT1^N/C^ (fig. S7B).

The Nucleoprotein is a critical multifunctional protein, which protects the viral RNA, interacts with and recruits host components to IBs, thereby aiding in IB formation and enabling viral genomic RNA replication within the IBs, among its several other functions (26–35). EHMTs were found to methylate the lysine residue/s of only the viral Nucleoprotein upon infection, inhibition of which led to reduction in levels of the Nucleoprotein. Since the viral Nucleoprotein has been implicated in almost every stage of the viral life cycle and EHMTs regulated its levels via methylation, it is prudent to believe that these enzymes facilitate viral replication. In combination with methylation of N protein, EHMTs might have additional host specific substrates that aid in viral replication which still needs to be identified. Overall, our study has explored an underappreciated role of a host methyltransferase in viral pathogenesis. Identification of such protein-protein interaction is not only critical for SeV biology but can provide mechanistic insights into possibly conserved mechanism of host pathogen interaction networks of other members of the order mononegavirales.

Post translational modifications (PTMs) on proteins generate diversity of a protein pool (65) and are responsible for regulating a large number of biological processes in both host and pathogen (66–70). RNA viruses encode for a handful of proteins that are multifunctional in nature. Thus, harnessing host enzymes responsible for PTMs to impart diversity to their proteins, thereby increasing the affinity of protein: protein interactions can be a powerful strategy for viruses to establish efficient replication. In support of this phenomenon, we found that EHMT1^N/C^ methylates the SeV Nucleoprotein. Association of EHMT1^N/C^ with N and methylation of N increased as the infection progressed. The timeline of increased methylation of N protein coincided with the formation of larger inclusion bodies and increased replication of SeV. Consistent with these mechanistic insights, we observed that inhibiting the catalytic activity of EHMT1^N/C^ using small molecule inhibitors highly reduced the abundance of larger IBs and decreased SeV replication. Prior to our study, the only report which demonstrated involvement of a lysine methyltransferases in RNA viral replication organelles is by Chen et al., wherein Smyd3 facilitated transcription of Ebola virus (35). However, whether Smyd3 mediates methylation of viral proteins is unknown. In the current study, we for the first time demonstrate lysine methylation as a PTM to regulate the biomolecular condensation in the context of host-pathogen interaction.

Viruses are known to recruit enzymes such as kinases to phosphorylate several of their proteins. For example, human T-cell lymphotropic virus protein Tax (71), VP30 of Ebola (72), Nucleoprotein of Influenza (68), etc are known to get phosphorylated, which eventually aids in activating viral gene transcription. A series of biochemical studies showed that GCN5 and PCAF were responsible for acetylation of the nucleoprotein in Influenza virus (73). In addition, studies on viral envelope proteins identified its glycosylation to facilitate viral entry to dampen the immune response of the host (74). To our knowledge, this is the first study where we report the involvement of a lysine methyltransferase and its methyltransferase activity being directly utilized by ssRNA virus in the formation/maturation of IBs (or Replication Organelles) for efficient replication. Given that IB formation is a conserved phenomenon among several RNA viruses, it would be interesting to determine the conservation of EHMT1 in the formation of IBs and replication of other related members of SeV, which are human pathogens. Overall, our work might have strong impact in understanding pathogenic RNA virus biology and targeting therapeutic strategies against viral replication.

In a high throughput study performed to determine the enzymatic substrates of EHMT1 (6) several extranuclear proteins, including several mitochondrial, ER and cell membrane specific proteins were detected (6). These observations were puzzling given the nuclear subcellular localization EHMT1 (8). Our findings of a distinct isoform of EHMT1 (EHMT1^N/C^), can be instrumental in studying EHMT1^N/C^ mediated non-histone, non-nuclear protein regulations and their complex interactomes. While we have identified how EHMT1^N/C^ plays a significant role in RNA viral infection, its expression in the cytoplasm independent of infection in multiple cell lines indicates its potential role in regulating several biological processes. Knockout of EHMT1 in mice results in embryonic lethality (1), haploinsufficiency manifests as a neurodevelopmental disease called Kleefstra Syndrome (75), Copy Number Variations were reported in Schizophrenia (76) and elevated expression was found in several cancers (77,78) and Alzheimer’s Disease (79). Until now, our understanding about these diseases emanated from the regulation of processes governed by nuclear EHMT1 but the identification of a novel nucleo- cytoplasmic form extends beyond viral pathogenesis and is expected to enhance our understanding about the role of EHMT1 in development and disease.

In conclusion our findings discovered a previously unidentified nucleo-cytoplamic form of EHMT1 and its role as a pro-viral factor at the interface of host-virus interactions. At mechanistic level, our work demonstrated lysine methylation as a post translation modification of SeV N protein to build large inclusion bodies which are necessary for efficient replication. These findings will not only open new avenues to address questions pertaining to host-pathogens interaction and therapeutic interventions but will also be instrumental in illuminating unknown functions of EHMT1^N/C^ in the context of development and disease.

## Materials and methods

### Cell culture

Neonatal human dermal fibroblasts were purchased from ScienCell (2310), Mouse Embryonic Fibroblasts (MEFs) were isolated from mouse embryos 13 d.p.c. HEK 293, BEAS-2B were a gift from Dr. Arjun Guha. All the cell lines and primary cells were cultured in Dulbecco’s Modified Essential Media (DMEM, high glucose, GlutaMAX (Gibco)), supplemented with 10% (v/v) heat-inactivated Fetal Bovine Serum (Gibco), and 1X Non-Essential Amino Acids (Gibco). The cells were incubated in a 37°C, 5% CO2 incubator. For passaging, cells were briefly treated with 1X TrypLE (Gibco).

### Viruses

Cytotune OSKM Sendai virus (A16517, Invitrogen) and EmGFP reporter Sendai Virus (A16519, Invitrogen) were part of the Cytotune -iPS 2.0 Sendai Reprogramming kit and EmGFP Sendai Fluorescence Reporter kit. Wild Type Sendai Virus (Z strain) was propagated in the allantoic sacs of 10-day old embryonated chicken eggs. Cells were infected with a MOI of 2.

### Immunocytochemistry

Cells were seeded on glass coverslips of 0.167mm thickness at 40% confluency. 24h post seeding, they were either transfected or infected; the cells were then fixed with 4% Paraformaldehyde (w/v) for 10min at RT. This was followed by 3 PBS washes and permeabilization with 0.5% TritonX-100 (v/v) in 1% BSA for 10min at RT. The cells were then washed with PBS and blocked with 5% BSA for 1h at RT. Primary antibodies diluted in 5% BSA at recommended concentrations were incubated overnight at 4°C. The cells were then washed with PBS thrice and the respective fluorophore conjugated alexa secondary antibodies diluted in 5% BSA were incubated for 40min at RT. After PBS washes, DAPI was added for 5min at RT. The coverslips were then washed with PBS, air-dried, and mounted on glass slides with Vectasheild mounting media.

For co-immunolabelling with EHMT1 and SeV, we sequentially incubated the cells first with Rb EHMT1 (Novus) antibody at 1:100 dilution overnight at4°C, Chk SeV (abcam) antibody was added on the following day at 1:2500 dilution. The samples were incubated overnight at 4°C, followed by PBS washes. The alexa secondary fluorophore conjugated antibodies were mixed at a dilution of goat-anti-rabbit 647 (1:200) and goat-anti-chicken 568 (1:1000). The rest of the protocol followed is as explained in the previous paragraph.

### Confocal microscopy and Image analysis

Confocal microscopic imaging was performed for the samples either on Olympus FV3000 or Leica SP8 confocal microscope. Representative images were acquired from several fields in each sample. Images were extracted, pseudo colours were assigned based on the secondary antibody, dye or label used; z-stacks of the maximum intensity projection were constructed using ImageJ.

Live cell imaging was performed on cells seeded in coverslip bottom confocal dishes using the 63X oil immersion objective on the Leica SP8 confocal microscope.

Quantitation of the number and size of IBs was performed on ImageJ, for which Z-stacked images were converted to 8-bit and grayscale. The threshold was adjusted to eliminate the background and an ROI was marked for each cell to measure the area of each IB within a cell.

For object-based co-localization analysis, Volocity image analysis software from Perkin Elmer was used. Z-stacked raw images were processed for noise reduction on the Leica LAX software before importing to Volocity. The populations were defined based on the channels for SeV and EHMT1 and an ROI was marked for each cell, from which the nucleus was excluded. Object size guide was set at 25um^3^ to separate touching objects and the colocalization was measured.

### Cloning and Transfection

The SeV Nucleoprotein and Phosphoprotein were amplified from the cDNA of EmGFP SeV infected cells. The Nucleoprotein was inserted into the MCS of mCherry_C1 plasmid (#632524, Addgene) at the ECoR1 and BamH1 sites using the SeV Nucleoprotein primers. SeV Phosphoprotein was inserted into the MCS of piRFP670-N1 (#45457, Addgene) at the BglII and KpnI sites using the SeV Phosphoprotein primers. EHMT1 full length was subcloned from the V5 tagged EHMT1 plasmid into the MCS of mCherry_C1 plasmid at the BglII and KpnI sites using the mCh_EHMT1_FL primers. EHMT1 guide RNA was targeted against the exon3 of EHMT1 gene; it was cloned into the pSpcas9(BB)-2A-GFP (PX458) (#48138, Addgene). For episomal reprogramming, pCXLE-hSK (#27078), pCXLE-hOCT3/4-shp53-F (#27077) and pCXLE-hUL (#27080) plasmids were a gift from Shinya Yamanaka (80). These plasmids were transfected in an equal ratio.

For the nuclear EHMT1 compensation experiments, HEKs at 40% confluency were transfected simultaneously with EH_132/spcas9 and mCh_EH1_FL plasmids. 36h post transfection, the cells were trypsinised and seeded for infection at 48h. The cells were then harvested 16h post infection with the WT SeV.

### RNA extraction and qRT-PCR

Harvested cells were resuspended in Trizol for total RNA extraction by the conventional method using chloroform for phase separation and Isopropanol for precipitation of RNA. Reverse transcription of RNA to cDNA was performed using the PrimeScript RT Reagent kit (RR037A, Takara). Quantitative RT-PCR was then performed on the cDNA using TB Green Premix Ex Taq II (RR820A, Takara), following the instructions manual.

### Nuclear cytoplasmic fractionation and Western Blotting

The cells were harvested by trypsinization, and the pellet was washed twice with PBS. The pellet was then resuspended in 0.1% NP40 in PBS, supplemented with 1X Protease Inhibitor Cocktail (PIC) and triturated about 10 times on ice. The samples were then incubated on ice for 1min and centrifuged at 1000g for 10min at 4°C. The resulting supernatant was the cytoplasmic fraction, which was aspirated into a fresh tube. The pellet was washed twice by resuspending in 0.1% NP-40 in PBS and centrifugation at 1000g for 10min at 4°C. The resulting pellet was then resuspended in RIPA buffer + 1X PIC, incubated on ice for 1h and vortexed at regular intervals. The samples were then centrifuged at 12,000g for 10min at RT to eliminate the debris, the resulting supernatant was the nuclear fraction.

Whole cell lysate was prepared by resuspending the cell pellet in RIPA buffer + 1X PIC, which was incubated on ice for 1h and vortexed at regular intervals.

The lysates and fractions were estimated for protein concentration by Bradford Assay. Required amounts of the protein were reduced and denatured by mixing with 1X NuPAGE Sample Reducing Agent (Thermofisher Scientific) and IX NuPAGE LDS Sample Buffer and heating at 70°C for 10min. The samples were then resolved on either 8% or 4% SDS-PAGE gel and immunoblotted with respective antibodies. The bands obtained by Western Blotting were quantified by ImageJ.

### Antibodies, Inhibitors and Reagents

EHMT1 (NBP1-77400, Novus Biologicals, Rabbit polyclonal) was used for ICC, EHMT1 (A3 01-642A, Thermofisher Scientific, Rabbit polyclonal) was used for ICC, WB, IP. Sendai virus (ab33988, abcam, Chicken polyclonal), Sendai virus (PD029, MBL, Rabbit polyclonal), EHMT2 (NBP2-13948, Novus Biologicals, Rabbit polyclonal), Ezh2 (D2C9, Cell Signalling Technology, Rabbit monoclonal), Oct4 (9B7, MAI-104, Invitrogen, Mouse monoclonal), Hsp70 (W27, sc-24, Santa Cruz, mouse monoclonal), Gapdh (G9545, Sigma, Rabbit polyclonal), panH3 (ab1791, abcam, Rabbit polyclonal), H3K9me2 (ab32521, abcam, Rabbit monoclonal), LaminB1 (ab16048, abcam, Rabbit polyclonal), mCherry (NBP1-96752, Novus Biologicals, mouse monoclonal), Methylated e-N Lysine (ICP0501, Immunechem, Rabbit polyclonal), Normal Rabbit IgG (12-370, Sigma-aldrich, Rabbit polyclonal), Normal Mouse IgG (12-371, Sigma-aldrich, Mouse polyclonal).

For ICC, Alexa Fluor conjugated secondary antibodies from Invitrogen were used: Goat anti-Rabbit 633 (A21071), Goat anti-Rabbit 568 (A11011), Goat anti-chicken 568 (A11041), Goat anti-mouse 488 (A11001), Goat anti-Rabbit 488 (A11008), Goat anti-mouse 568 (A11004). HRP-conjugated secondary antibodies were used for Western Blotting: Goat anti-Rabbit (1706515, Biorad), Goat anti-mouse (1706516, Biorad). BIX-01294 (B9311, Sigma), UNC0642 (SML1037, Sigma). Dynabeads Protein A (10001D, ThermoFisher Scientific), Dynabeads Protein G (10003D, ThermoFisher Scientific).

### Cell infection, treatment, and transfection

Cells were infected with WT SeV, Cytotune SeV or EmGFP SeV at a Multiplicity of Infection of 2 in culture media. Cells were treated with 3μM of BIX-01294 or 3μm of UNC-0642, which was replenished every 24h. Plasmids were transfected in HEK by using Xfect Transfection Reagent (631318, Takara) by following the user manual.

### T7 endonuclease assay

Genomic DNA was isolated and 550bp region of the exon 3 was amplified, with about 250 bases flanking on either side of the gRNA targeted region.

### Immunoprecipitation and RIP

Dynabeads protein A or G were washed twice with 0.1% BSA in 1M Potassium phosphate buffer. To this, 2ug of the respective antibody diluted in 0.5M Potassium phosphate buffer was added and incubated on a rotor for 2h at RT. The unbound antibody was then washed with Potassium Phosphate buffer and the protein lysate, or the cytoplasmic fraction was added and incubated ON at 4°C. (For Immunoprecipitation, the cytoplasmic fraction was mixed 1: 1 with RIPA buffer). On the following day, the unbound fraction was removed by washing with IP-100 and 1X Flag buffer. The Immunoprecipitated protein was then eluted by incubating in 2X reducing agent and protein loading dye at 70°C for 10min.

### RNA Immunoprecipitation

The cells were washed with PBS in the culture dish and UV-crosslinked at 450 mJ/cm^2^ before proceeding with Nuclear cytoplasmic fractionation or cell lysis. About 750- 1000ug of protein was added to the antibody bound dyna beads and incubated ON at 4°C. On the following day, the unbound fraction was removed by washing with IP-100 or 1X Flag buffer. Immunoprecipitated RNA was eluted from the beads by treating with 5mg/ml ProteinaseK for 15min at 37°C; the sample was then incubated with trizol for RNA extraction by the chloroform/isopropanol method.

### Statistical analysis

Graphpad Prism 8 was used to perform all the statistical analyses. For analysis of two groups, paired or unpaired t-test was used according to the experiment. For comparison between three or more groups, One-way ANOVA or Brown-Forsythe and Welch ANOVA was used. Details of N (replicate size), n (sample number) and the tests applied have been mentioned in the respective figure legends. p-value: 0.1234 (ns), 0.0332 (*), 0.0021 (**), 0.0002 (***), <0.0001 (****).

## Acknowledgements

This project was supported by funds from DBT/Wellcome Trust India Alliance Intermediate Fellowship (Grant number # 500220-Z-11-Z) to SR, funds from CSIR-IGIB (OLP 1153) to SR and inStem Core funds to PKV. K.K.B is supported by CSIR-JRF/SRF fellowship. NRS received support from the Ramalingaswami Re-entry Fellowship from the Department of Biotechnology (DBT), Ministry of Science and Technology, Government of India (BT/RLF/Re-entry/40/2018). We thank Prof. Colin Jamora and Dr. Dasaradhi Palakodeti for their critical and constructive feedback on the manuscript. We thank Dr. Sheetal Gandotra for the Volocity Image analysis software and helpful discussions. The Central Imaging and Flow Cytometry Facility (CIFF) at InStem and NCBS and Confocal microscopy facility at CSIR-IGIB supported the confocal microscopic imaging.

## Author Contributions

S.R. and K.K.B. conceptualized the project and designed the experiments. K.K.B. performed all the experiments and analysed the data. K.K.B. and S.R. wrote the manuscript. D.P.S/N.R.S. provided wild type Sendai Virus, SeV antibody and provided intellectual inputs on the project. A.M generated the construct for EHMT1 truncations. P.K.V. provided funds and intellectual inputs on the project.

## Competing interests

The authors declare no competing interests.

## Supplementary Information

**Supplementary figure 1:**
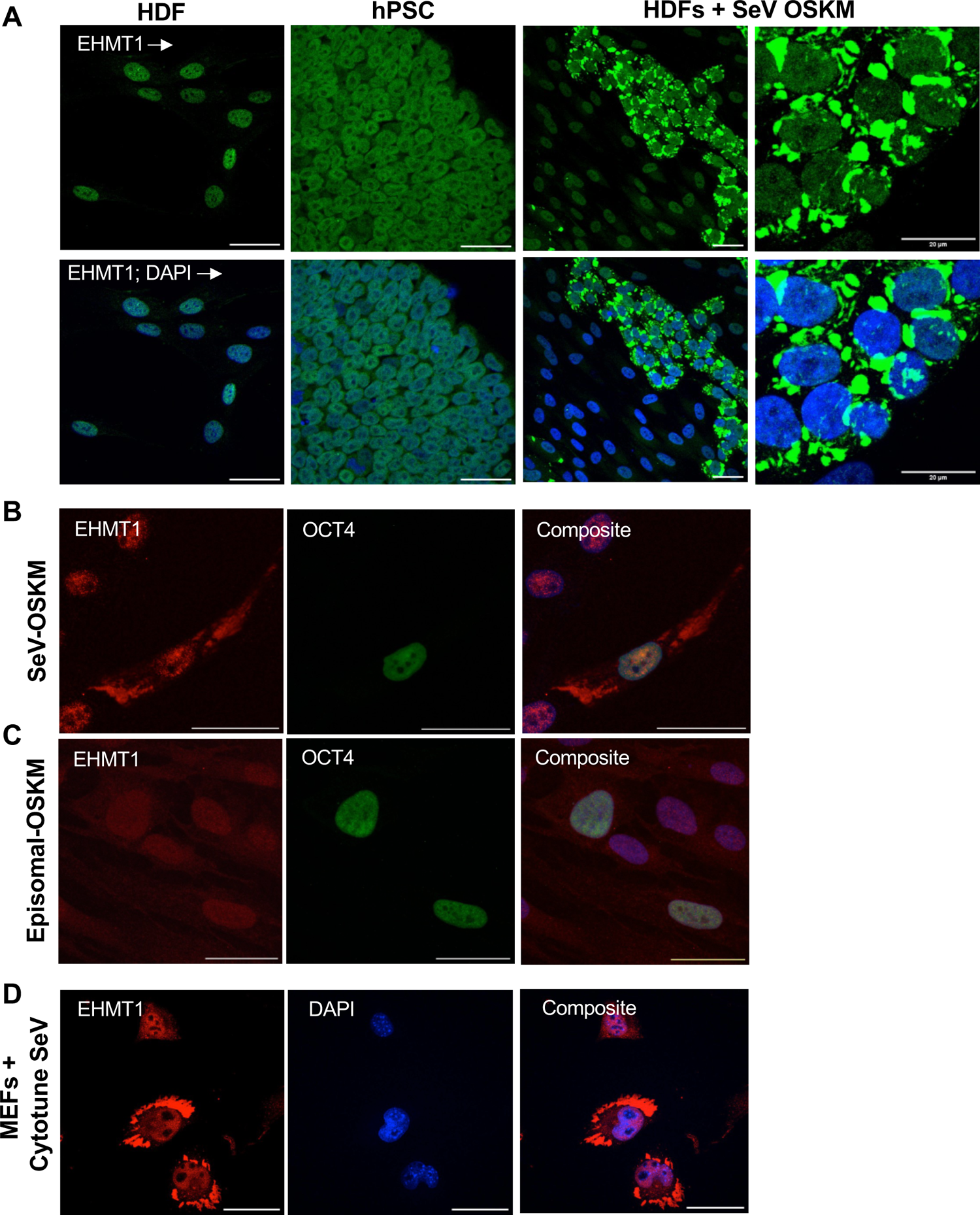
Transduction of Sendai OSKM virus in fibroblasts resulted in formation of EHMT1 condensates in the cytoplasm. Confocal microscopic images of (A) Various stages of fibroblasts undergoing reprogramming induced by the ectopic expression of OSKM delivered via SeV, immunolabelled with EHMT1 (green). (B) Fibroblasts transduced with OSKM via SeV, immunolabelled for EHMT 1 (red) and Oct4 (green). (C) Fibroblasts transfected with episomal plasmids expressing OSKM, immunolabelled with EHMT1 (red) and Oct4 (green). (D) Mouse Embryonic Fibroblasts (MEFs) infected with SeV, immunolabelled with EHMT1 (red). Composite of images are with DAPI (blue) stained nuclei. Scale bar - 40μm.

**Supplementary figure 2:**
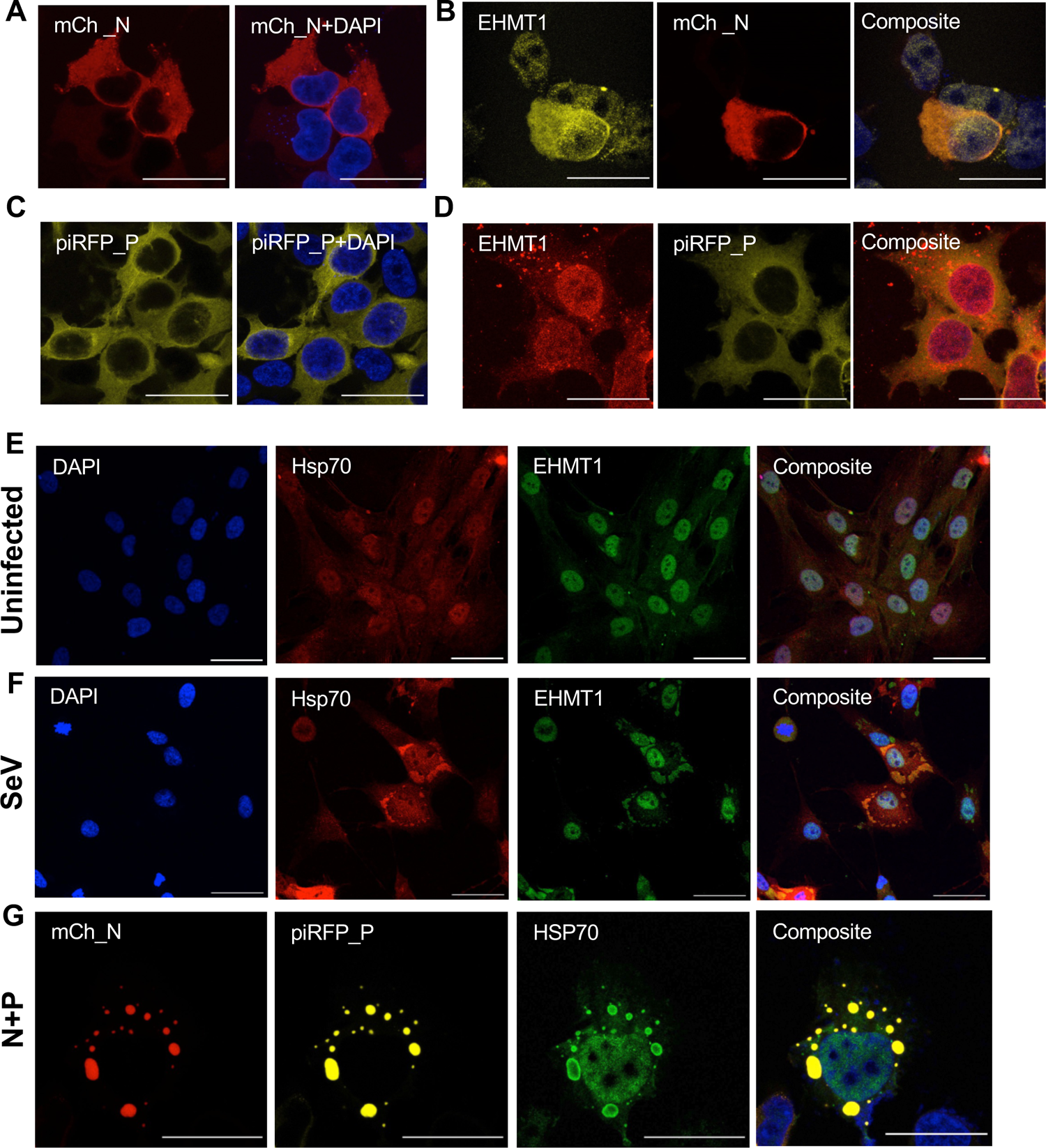
Expression of SeV N and P proteins independent of SeV infection. Confocal microscopic images of HEKs transfected with (A) mCh_N (red) and (C) piRFP_P (yellow), (B) HEKs transfected with mCh_N (red) and immunolabelled with EHMT1 (yellow), (D) HEKs transfected with piRFP_P (yellow) and immunolabelled with EHMT1 (red). (E) Fibroblast uninfected and (F) infected with SeV co-immunolabelled with Hsp70 (red) and EHMT1 (green). (G) HEKs co­transfected with mCh_N (red) and piRFP_P (yellow) in a 1:1 ratio, immunolabelled with Hsp70 (green). Composite of all images are with DAPI (blue) stained nuclei. Scale bar - 40μm.

**Supplementary figure 3:**
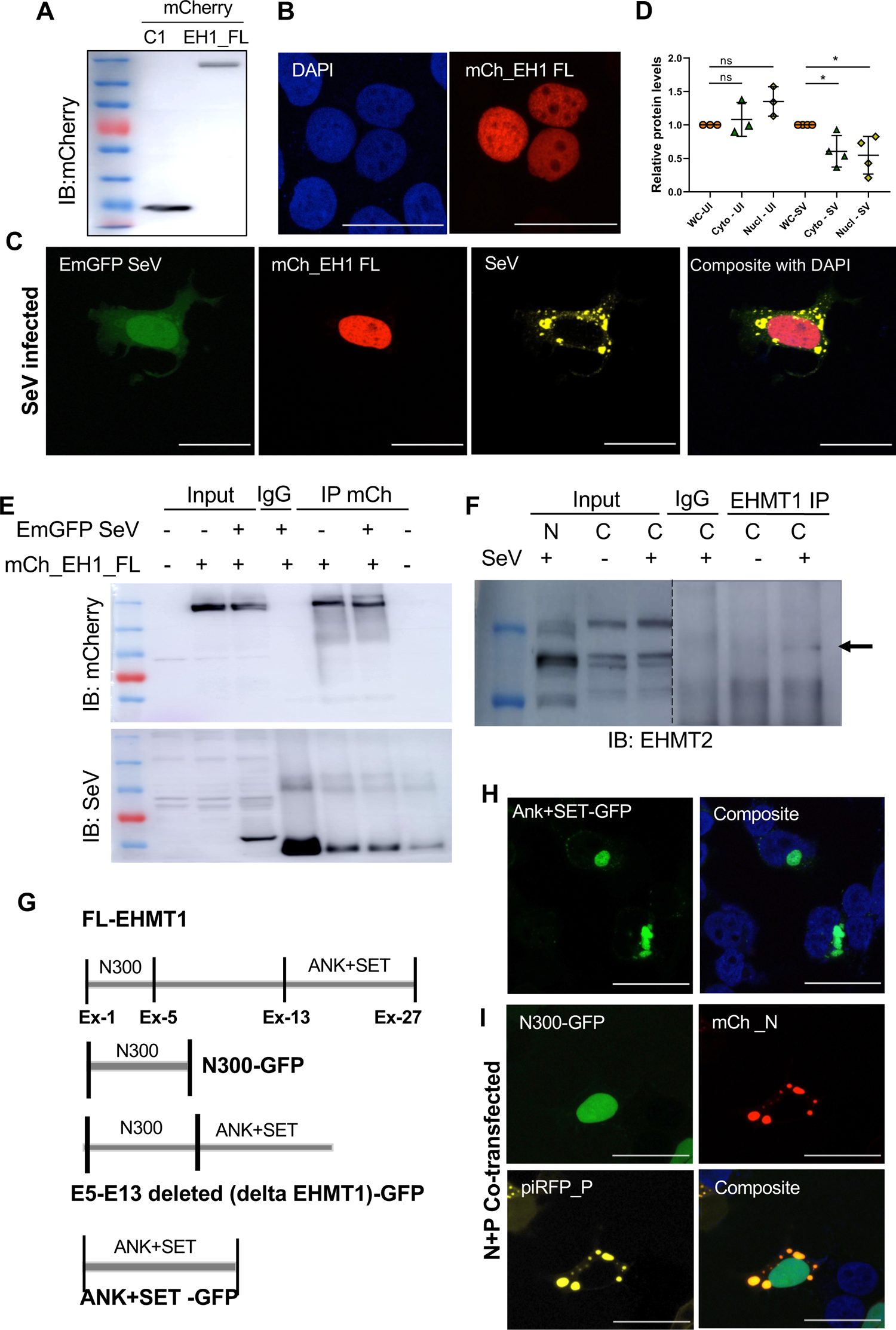
Distinct cytoplasmic form of EHMT1 localises with SeV IBs. (A) Whole cell lysate of HEKs transfected with mCh_C1 and mCh_EH1_FL, were Western blotted and probed with mCherry. Confocal microscopic images of HEK transfected with (B) mCh_EH1_FL (red), (C) mCh_EH1_FL (red), infected with EmGFP SeV (green) and immunolabelled with SeV (yellow). Nuclei stained blue with DAPI. (D) Graph representing relative protein levels of EHMT 1 quantified from Western blots of Whole cell, nuclear and cytoplasmic fractions at 24h p.i.. Cytoplasmic and nuclear protein levels were normalised against Whole cell (n ≥ 3 replicates, one-way ANOVA, p-value: 0.1234 (ns), 0.0332 (*)). (E) mCherry IP from whole cell lysate of HEK transfected with mCh_EH1_FL and infected with EmGFP SeV; the elute was Western blotted and probed with mCherry and SeV, depicting no association between mCh_EHMT1_FL and SeV proteins. (F) EHMT1 IP from the cytoplasmic fraction of uninfected/SeV infected cells; the elute was western blotted and probed with EHMT2, depicting an interaction between EHMT1 and EHMT2. (G) Schematic representation of EHMT1 truncated sequences cloned into EGFP plasmid. (H) Confocal microscopic images of HEK transfected with Ank+SET_EGFP. (I) Confocal microscopic images of HEK transfected with N300-GFP, co­transfected with mCh_N + piRFP_P, composite images are with DAPI (blue) stained nuclei. Scale bar - 40μm.

**Supplementary figure 4:**
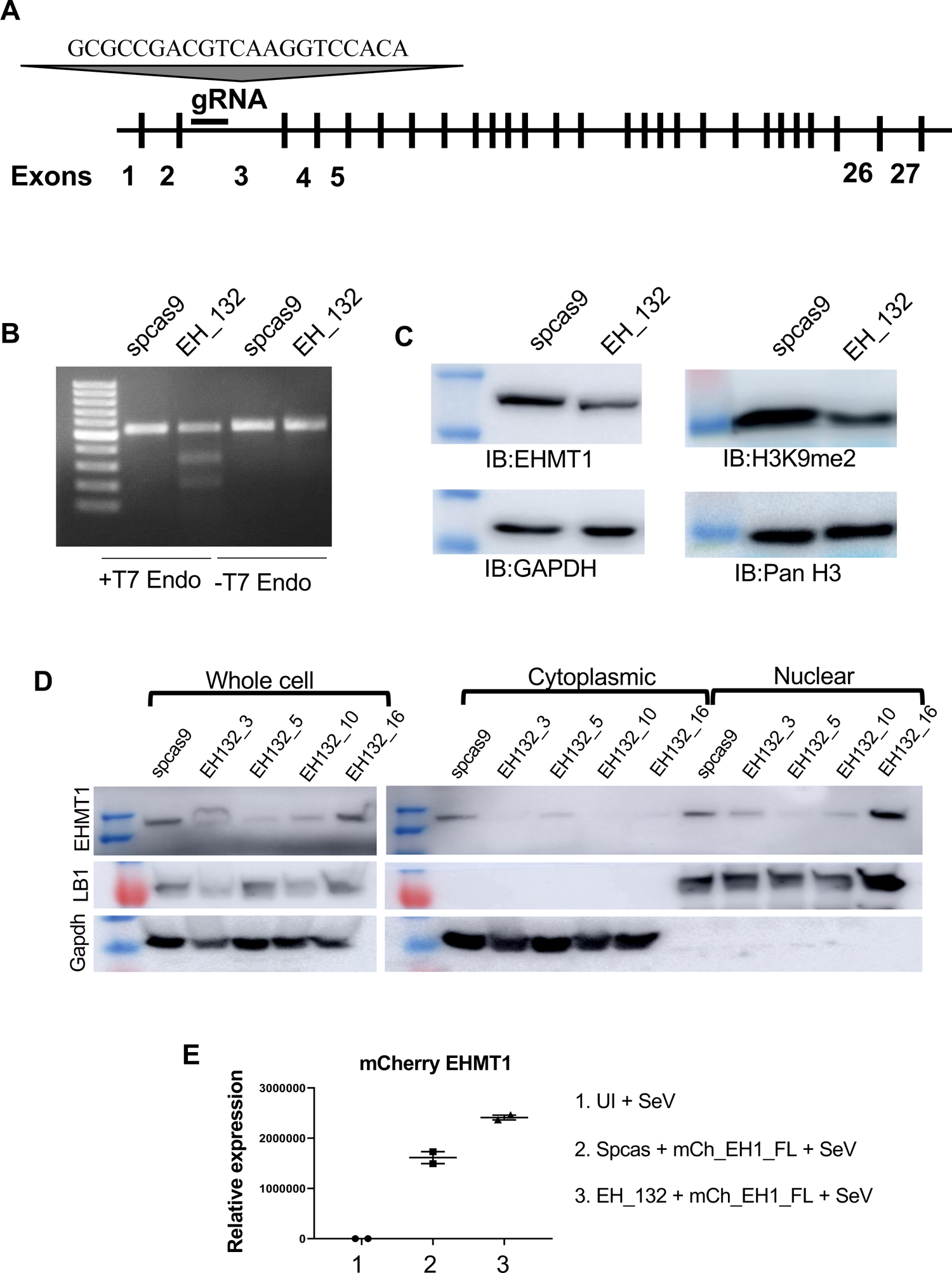
Depletion of EHMT1 by CRISPR/cas9. (A) Schematic and sequence of EH_132 guide RNA targeting exon 3 of EHMT1 via CRISPR/cas9 method of gene editing. (B) Agarose gel electrophoresis of the products from T7 endonuclease assay demonstrating gene editing as seen by digestion into three products in EH_132 mutated sample. (C) Western blotting of the spcas9 control and EH_132 lysates immunoblotted with EHMT1, Gapdh, H3K9me2 and panH3. (D) Western blotting of the Whole cell, nuclear and cytoplasmic fractions of single cell clones of EH_132, probed with EHMT 1, LaminB1 and GAPDH. (E) Graph representing the relative mRNA levels of mCh_EH1_FL determined by qRT-PCR.

**Supplementary figure 5:**
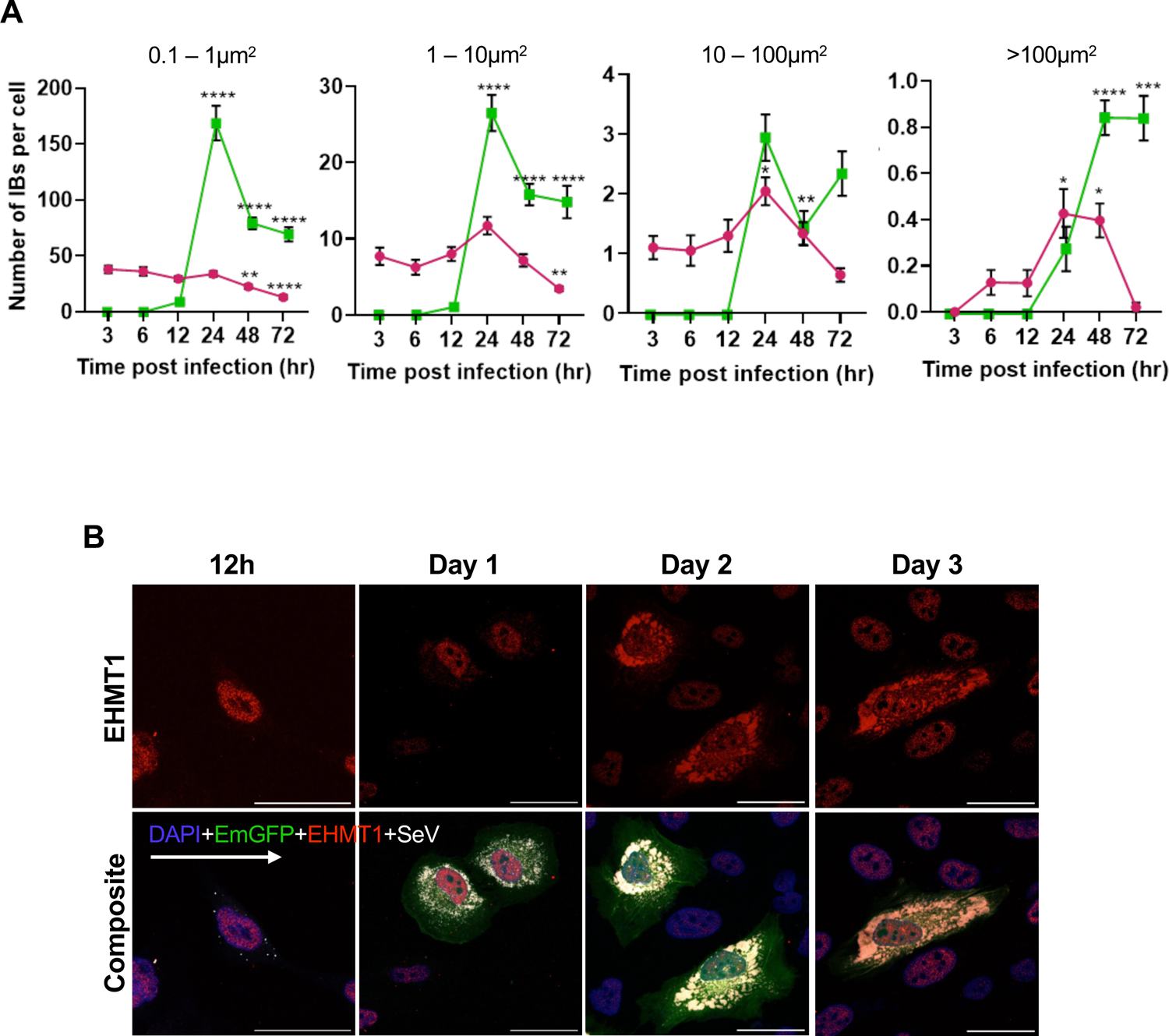
EHMT1^N/C^ differentially localises to large IBs. (A) Graphs representing the mean number of IBs (y-axis) in each subpopulation for EmGFP (green line) and WT SeV (pink line) infected cells. Data are ± S.E.M. (n>25 cells, Brown-Forsythe and Welch ANOVA tests), p-value: 0.1234 (ns), 0.0332 (*), 0.0021 (**), 0.0002 (***), <0.0001 (****). (B) Confocal microscopic images of BEAS-2B cells infected with EmGFP SeV (green) immunolabelled with EHMT1 (red) and SeV (gray) at indicated time points post infection, Composite images of all channels are with DAPI (blue) stained nuclei.

**Supplementary figure 6:**
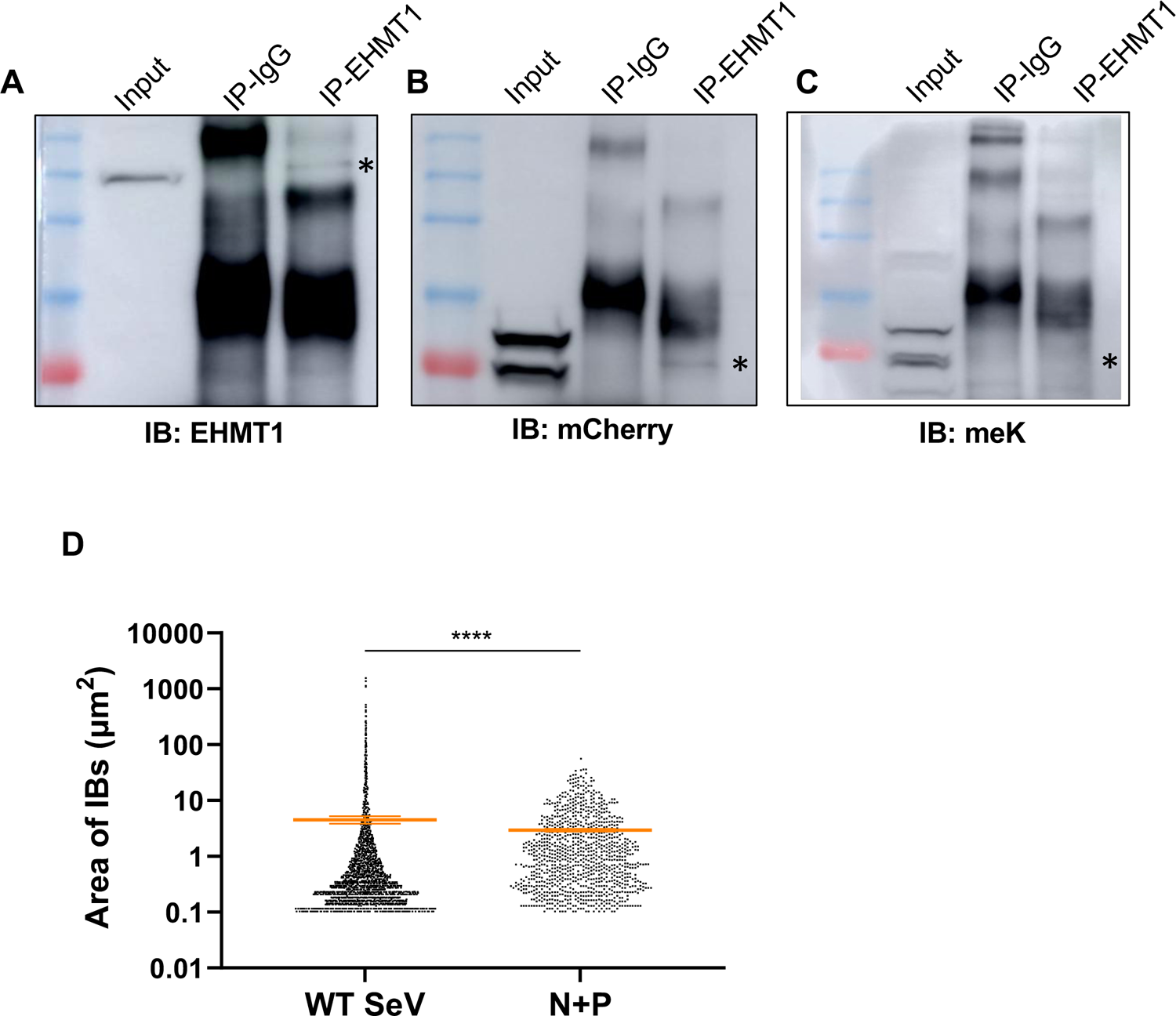
EHMT1^N/C^ mediated methylation of SeV Nucleoprotein occurs only during infection. (A) to (C) HEK were co-transfected with mCh_N and piRFP_P, EHMT1^N/C^ was immunoprecipitated from the cytoplasmic fraction 24h post transfection. IP elute was Western blotted and probed with (A) EHMT 1, (B) mCherry and (C) meK. (D) Graph representing the distribution of area of IBs formed in WT SeV infected cells and N+P co-transfected cells. Data are ± S.E.M. (N=3 replicates, n>100 cells, Unpaired t-test, <0.0001 (****)).

**Supplementary figure 7:**
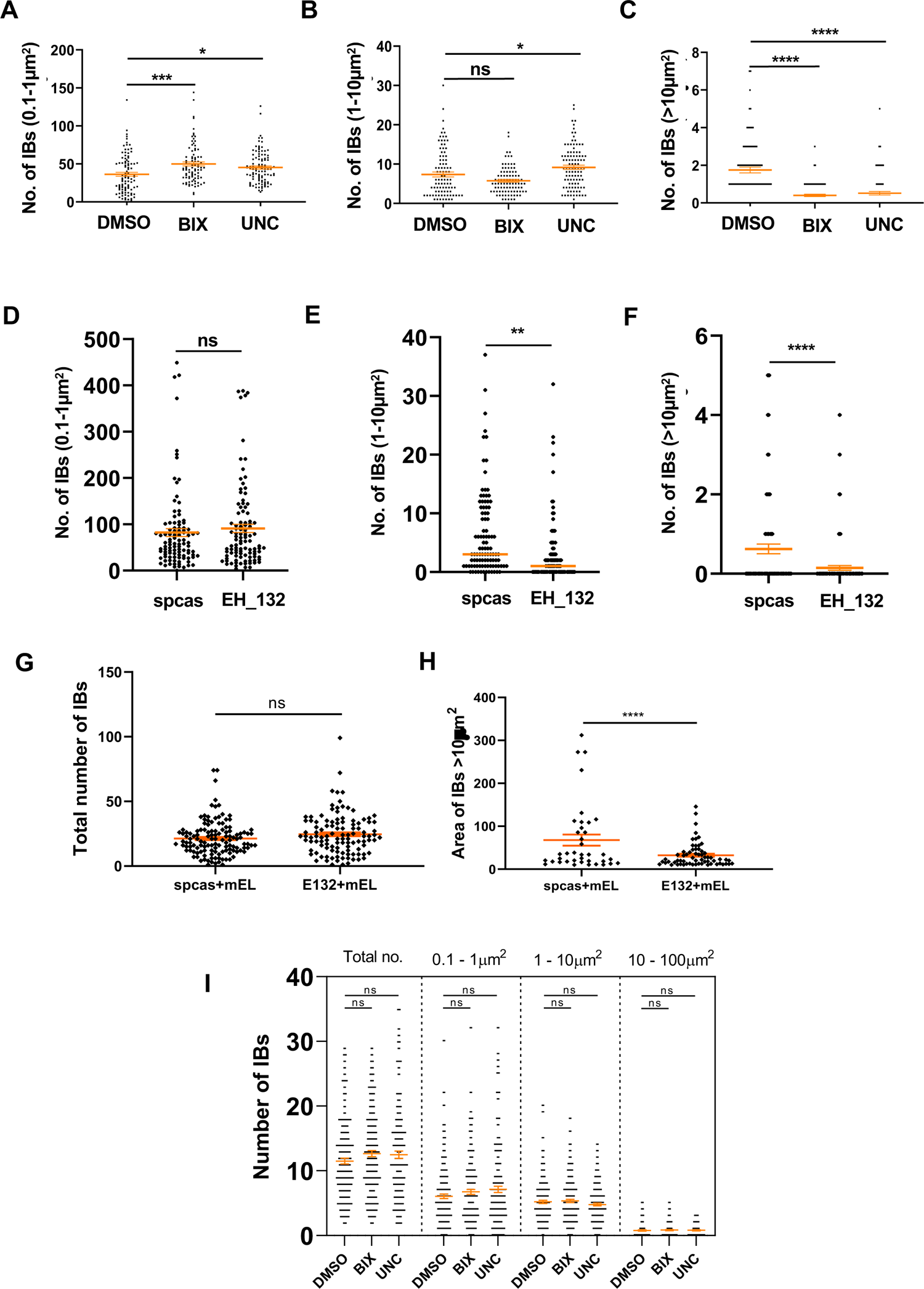
Enzymatic activity of EHMT1^N/C^ is needed for the growth of IBs upon infection. BEAS-2B cells were infected with WT SeV and treated with 3 μm of BIX/UNC simultaneously. The cells were fixed and immunolabelled with SeV antibody 16h p.i., the number of IBs and their area was quantified by ImageJ. Graphs representing number of IBs in (A) 0.1 - 1pm^2^ category, (B) 1-10 pm^2^ category, (C) >10 pm^2^ category. HEK293 cells were transfected with EH_132 plasmid to deplete the levels of EHMT1 and infected with WT SeV, cells were immunolabelled with SeV 16h p.i., number and area of IBs was quantified by ImageJ. Graphs representing number of IBs in (D) 0.1 - 1μm^2^ category, (E) 1-10 pm^2^ category, (F) >10 pm^2^ category. HEK293 were transfected with spcas9 empty plasmid or EH_132, and mCh_EH1_FL (mEL); the cells were then infected with WT SeV and immunolabelled for SeV 16h p.i. The number and area of IBs we quantified by ImageJ. (G) Graph representing the total number of IBs, (H) Graph representing the area of IBs >10pm^2^. (I) HEKs were co-transfected with mCh_N and piRFP_P, treated with DMSO, BIX or UNC and fixed at 24h to analyse IB formation by confocal microscopy followed by image analysis via ImageJ. Graph representing the total number of IBs and the number of IBs in each subpopulation. Data are ± S.E.M. (N=5 replicates, n>300 cells, Ordinary one-way ANOVA, 0.1234 (ns))

### Primer sequences used for cloning

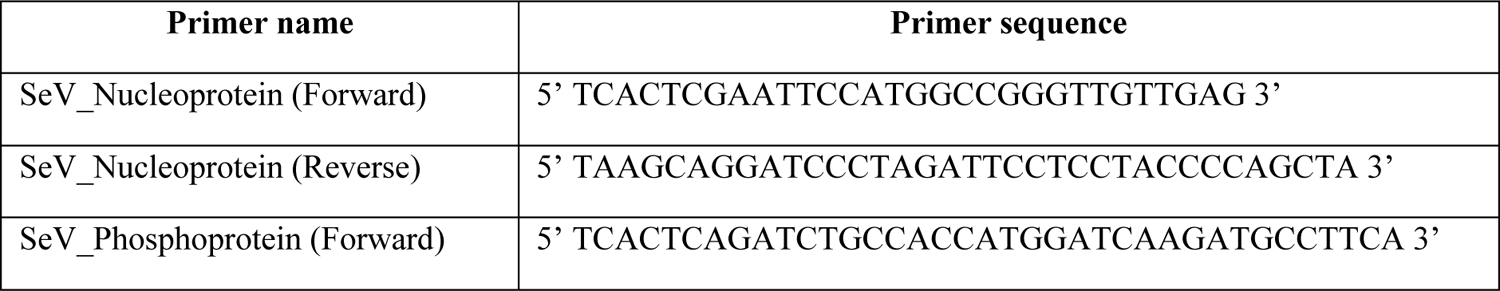
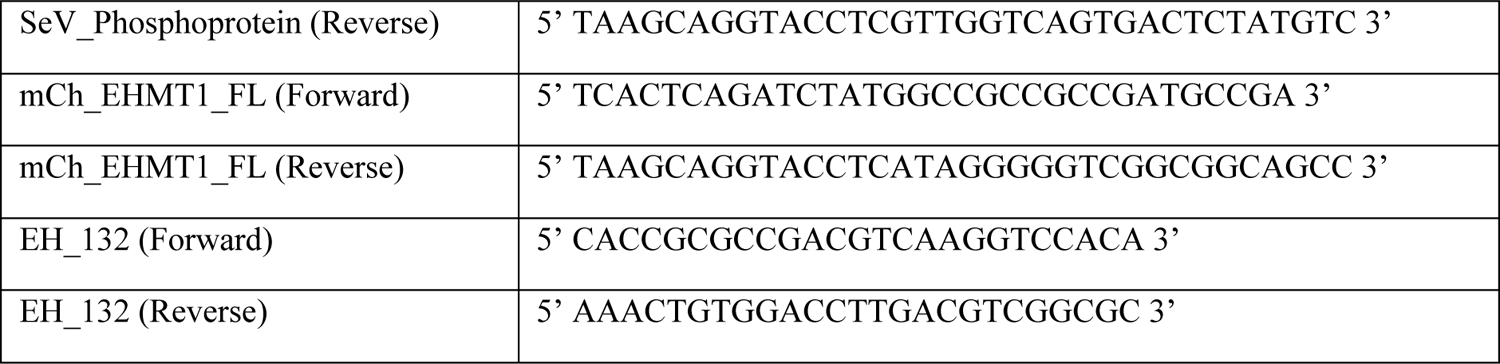

### Primer sequences used for qRT-PCR

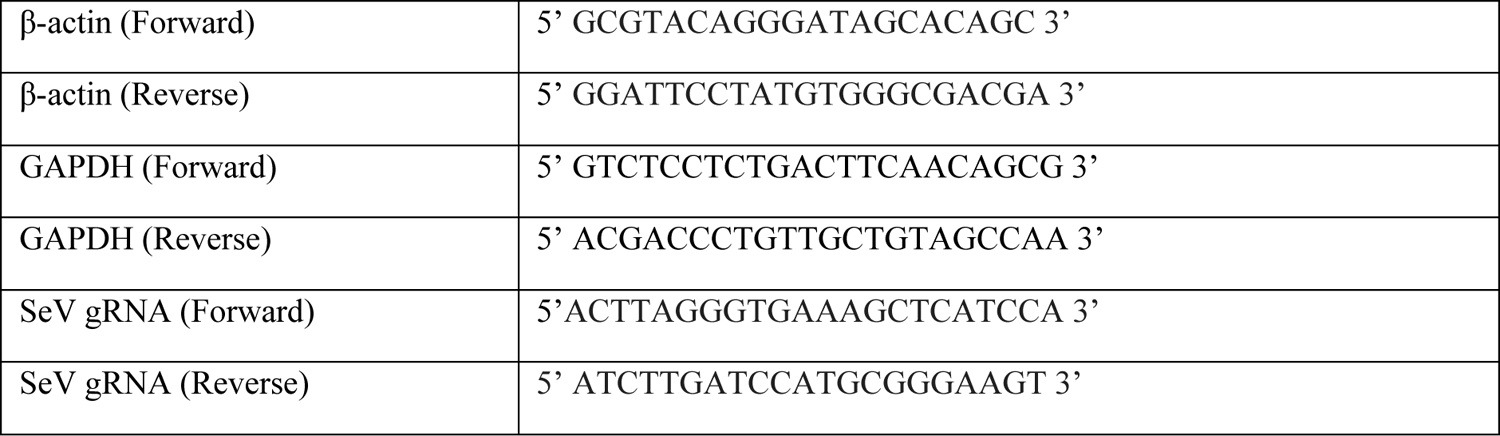

## Notes

### Competing Interest Statement

The authors have declared no competing interest.

